# Moesin controls cell-cell fusion and osteoclast function

**DOI:** 10.1101/2024.05.13.593799

**Authors:** Ophélie Dufrancais, Perrine Verdys, Marianna Plozza, Arnaud Métais, Marie Juzans, Thibaut Sanchez, Martin Bergert, Julia Halper, Christopher J Panebianco, Rémi Mascarau, Rémi Gence, Gaëlle Arnaud, Myriam Ben Neji, Isabelle Maridonneau-Parini, Véronique Le Cabec, Joel D Boerckel, Nathan J Pavlos, Alba Diz-Muñoz, Frédéric Lagarrigue, Claudine Blin-Wakkach, Sébastien Carréno, Renaud Poincloux, Janis K Burkhardt, Brigitte Raynaud-Messina, Christel Vérollet

## Abstract

Cell-cell fusion is an evolutionarily conserved process that is essential for many functions, including fertilisation and the formation of placenta, muscle and osteoclasts, multinucleated cells that are unique in their ability to resorb bone. The mechanisms of osteoclast multinucleation involve dynamic interactions between the actin cytoskeleton and the plasma membrane that are still poorly characterized. Here, we found that moesin, a cytoskeletal linker protein member of the Ezrin/Radixin/Moesin (ERM) protein family, is activated during osteoclast maturation and plays an instrumental role in both osteoclast fusion and function. In mouse and human osteoclast precursors, moesin inhibition favors their ability to fuse into multinucleated osteoclasts. Accordingly, we demonstrated that moesin depletion decreases membrane-to-cortex attachment and enhances the formation of tunneling nanotubes (TNTs), F-actin-based intercellular bridges that we reveal here to trigger cell-cell fusion. Moesin also controls HIV-1- and inflammation-induced cell fusion. In addition, moesin regulates the formation of the sealing zone, the adhesive structure determining osteoclast bone resorption area, and thus controls bone degradation, via a β3-integrin/RhoA/SLK pathway. Supporting our results, moesin*-*deficient mice present a reduced density of trabecular bones and increased osteoclast abundance and activity. These findings provide a better understanding of the regulation of cell-cell fusion and osteoclast biology, opening new opportunities to specifically target osteoclast activity in bone disease therapy.

## INTRODUCTION

Cell-cell fusion is a biological process where two or more cells combine to form a single cell with a shared cytoplasm and a single, continuous plasma membrane^1^. This phenomenon plays a crucial role in various physiological processes, including fertilization and in the development of certain tissues, organs and specialized cells, such as multinucleated bone-resorbing osteoclasts^2,3^.

Multinucleated osteoclasts are the exclusive bone-resorbing cells essential for bone homeostasis, which also have immune functions ^4^. They differentiate through the concerted action of macrophage-colony-stimulating factor (M-CSF) and receptor activator of NF-κB ligand (RANKL) ^5^. Postnatal maintenance of osteoclasts is mediated by acquisition of new nuclei from circulating blood cells that migrate towards bones and fuse with multinucleated osteoclasts in contact with the bone matrix ^6–9^. Although *in vitro* studies suggest that the fate of osteoclasts is to die by apoptosis ^5^, multinucleated osteoclasts can also undergo fission producing smaller cells, called osteomorphs, that can fuse again to form new osteoclasts ^8^. Mature osteoclasts contain up to around 20 nuclei *in vivo* ^10^ and control of osteoclast fusion appears crucial for bone resorption as the multinucleation degree and the osteoclast size are most often correlated with osteolysis efficiency ^3,11,12^. Osteoclast fusion is a highly coordinated process that involves the migration of precursor cells towards one another, establishment of a fusion-competent status and initiation of cell-to-cell contacts, cytoskeletal reorganization, and finally fusion of their membranes ^13^. Upon attachment to bone, multinucleated mature osteoclasts form an F-actin rich structure crucial for bone resorptive activity called the sealing zone. This bone-anchored adhesion structure demarcates the area of bone resorption from the rest of the environment and consists of a complex assembly of podosomes ^14–18^. Each of these steps of osteoclastogenesis involves rearrangements of the actin cytoskeleton and its interactions with the plasma membrane, but, the precise mechanisms and sequence of events still remain poorly understood ^1,3^. As an example, osteoclasts can form tunneling nanotubes (TNTs) ^3,19–22^, F-actin-containing intercellular membranous channels representing a direct way of communication ^23,24^, but their characteristics and the molecular actors involved in their formation, stability or function are poorly defined ^20–22^.

Ezrin, Radixin and Moesin (ERM) proteins compose a family of proteins linking the actin cytoskeleton with the plasma membrane. Thereby, they regulate various fundamental cellular processes that involve the remodeling of the cell cortex such as cell division and cell migration ^25–29^. Phosphorylation of a conserved threonine residue in their C-terminal actin-binding domain activates them by stabilizing their open-active conformation, thereby favoring actin attachment to the plasma membrane. This phosphorylation is mediated by several kinases including the Rho kinase ROCK, the isoenzyme protein kinase C (PKC) and the Ste20-like l-kinase (SLK) ^30^. ERM proteins are widely expressed in a developmental and tissue-specific manner, with distinct as well as overlapping distribution patterns and functions ^27,31^. In leukocytes, ezrin and moesin are predominantly expressed ^32,33^, and they have unique or redundant functions in cell adhesion, activation and migration, as well as in the formation of the phagocytic cup and the immune synapse ^30,33–35^. Moesin-deficient (*Msn*-/-) mice exhibit T, B and NK cell defects, underscoring an important role for moesin in lymphocyte homeostasis ^32,35,36^. In the context of HIV-1 infection, ezrin, and to a lesser extent moesin, are involved in fusion-dependent virus entry and replication ^37–39^, and in the regulation of the virological synapse and virus-induced cell-cell fusion ^40,41^.

Although cell-cell fusion and osteoclastogenesis involve dynamic interactions between the actin cytoskeleton and the plasma membrane, until now, the role of cortex rigidity and ERM proteins in these processes have not been investigated. Here, we demonstrate that moesin is the major activated ERM in osteoclasts and that its depletion promotes *(i)* the fusion of osteoclast precursors, which correlates with the efficiency of TNT formation and reduced membrane-to-cortex attachment, and *(ii)* the formation of the sealing zones in mature osteoclasts, and consequently bone degradation. In osteoclasts, ERM activation is dependent on the β3-integrin/RhoA/SLK pathway. Importantly and consistently with our in vitro results, we report that moesin-deficient mice exhibit an osteopenic phenotype.

## RESULTS

### TNTs are essential for the osteoclast fusion process

During early stages of osteoclast formation, precursors form abundant TNT-like structures prior to cell-cell fusion ^3,22,24^. Here, to directly test the implication of TNTs in osteoclast fusion per se, we used two complementary osteoclast models: (i) osteoclasts derived from human blood monocytes (hOCs) and (ii) murine osteoclasts (derived from an immortalized myeloid cell line, mOCs) (**Figure S1A**) ^42,43^. In both models, F-actin staining showed the presence of podosomes (F-actin dots) but also of TNT-like structures at early stages (Day 3) of differentiation, whereas zipper-like F-actin structures, as described between osteoclasts ^44,45^, were more apparent during the later stages between adjoining multinucleated cells (**Figure 1A-B**). According to the definition of TNTs ^22–24,46,47^, we quantified TNTs as F-actin-positive structures that connect at least two cells and that do not adhere to the glass coverslip. Thick TNTs were classified based on their diameter (≥2 μm) and the presence of microtubules, versus thin TNTs, which were <2 μm and devoid of microtubules (**Figure 1C** and **S1B**) ^48^. We noticed that thick TNTs were usually positioned higher with respect to the substrate than the thin ones. The two types of TNTs were observed throughout the early stages of cell fusion (**Figure 1C**, **S1B, movies 1&2**). As osteoclast maturation progressed, the percentage of cells forming thick TNTs decreased, whereas those forming thin TNTs was unchanged (**Figure 1C**). Using live imaging in hOCs (**Figure 1D**, **S1C** and **movies 3-5**), we showed that the contact of a cell emitting a TNT-like structure with its cell partner and fusion of their cytoplasms took place within 90 min. Together, these results demonstrate that TNTs participate in the cell-cell fusion process and suggest that thick TNTs are preferentially required for osteoclast fusion.

**Figure 1:**
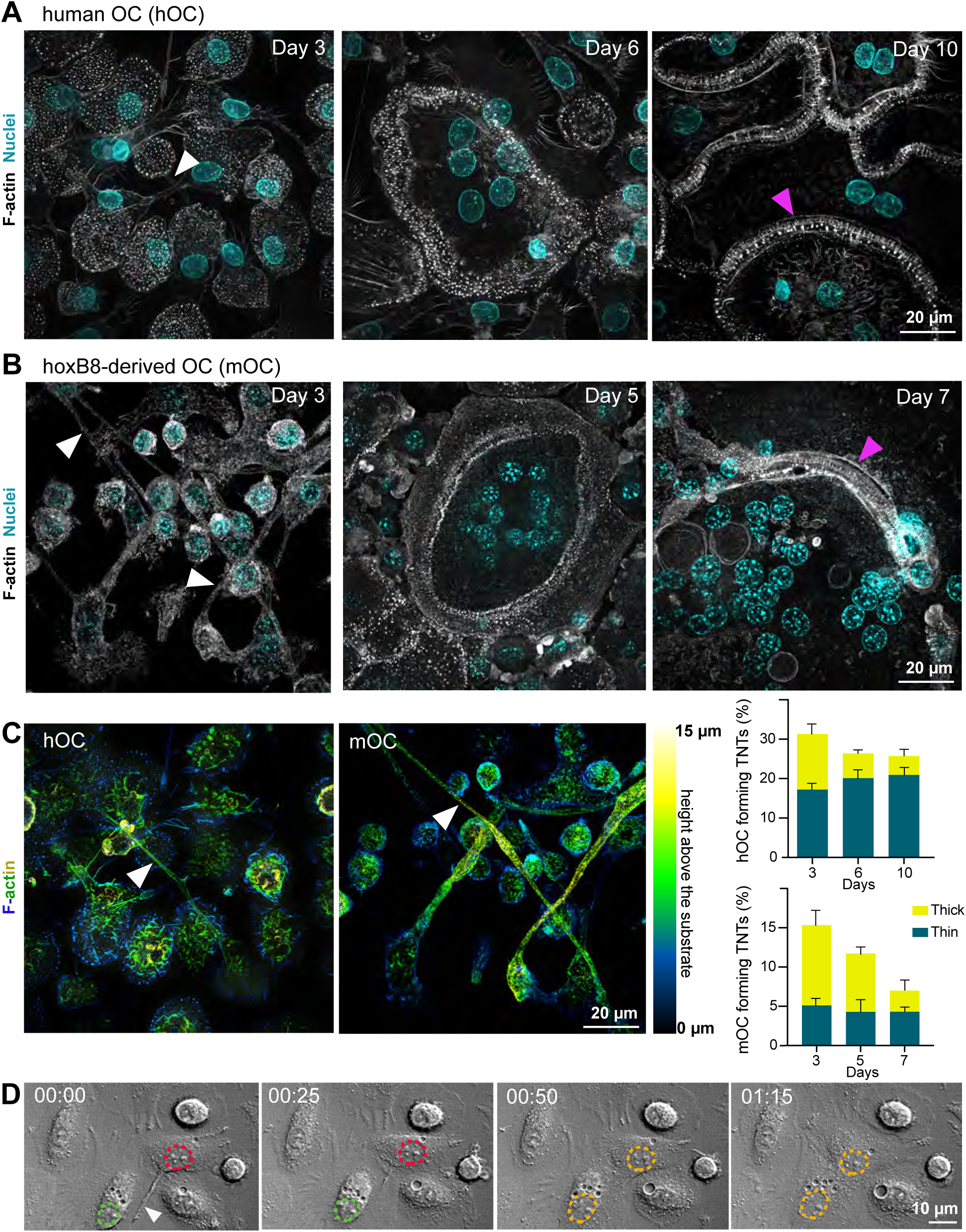
Tunneling nanotubes (TNTs) participate in the fusion of osteoclast precursors. **A.** Human monocytes isolated from blood were differentiated into osteoclasts (hOC) and analyzed at day 3, 6 and 10. Representative super-resolution microscopy images: F-actin (phalloidin, white) and nuclei (DAPI, cyan). Scale bar, 20 µm. **B.** Same experiment as in A with osteoclasts derived from the murine *HoxB8* immortalized cell line (mOC) at day 3, 5 and 7. White arrowheads show TNTs and pink arrowheads show zipper-like structures. **C.** Left panels: super-resolution microscopy images of TNTs with colored-coded Z-stack of F-actin (phalloidin) staining of 3 day-hOC or 3 day-mOC from 0 µm (substrate, dark blue) to 15 µm (yellow). Scale bar, 20 µm. See also **Movies 1&2**. Right panels: quantification of the percentage of cells forming thick and thin TNTs in hOCs and mOCs after immunofluorescence analysis (see methods), from one representative differentiation out of 3. n>250 cells per condition, means ± SEM are shown. **D.** Brightfield confocal images from a time-lapse movie of hOCs fusing from a TNT (hour:min). See **Movie 3**. Dashed green and red lines delineate the nuclei before cell fusion and dashed orange lines after fusion. Arrowhead shows a TNT-like protrusion. Scale bar, 10 µm.

### Moesin activation controls cell-cell fusion in several contexts

ERM proteins link the actin cytoskeleton to the plasma membrane and thereby regulate the formation of F-actin-based structures ^27^. We thus investigated the potential contribution(s) of ERM proteins during cell-cell fusion of osteoclasts. First, we confirmed that all three proteins ERM were expressed throughout mOC and hOC differentiation, confirming previous observations ^49,50^ (**Figure S2A-B**). Interestingly, we observed a strong increase in ERM activation status, as measured by the level of ERM phosphorylation (P-ERM) (**Figure 2A-B**), which peaked at Day 5 (mOC)/Day 6 (hOC), coinciding with the appearance of multinucleated osteoclasts (see **Figure 1A-B)**. To evaluate the function of ERM proteins, we engineered the individual knock-out (KO) of ezrin, radixin or moesin in mOCs. In each individual ERM KO, we did not observe compensation from the other ERM proteins (**Figure S2C**). While no difference in the cell-cell fusion was observed in the absence of either ezrin or radixin compared to controls (**Figure S2D**), deletion of moesin resulted in premature fusion of osteoclast precursors (**Figure 2C** and **movie 6**), leading to a significant increase in the fusion index, in the area occupied by osteoclasts, and in the number of nuclei per multinucleated cell (**Figure 2D-E and S2E**). We also observed a higher number of cells expressing the osteoclast maturation marker β3-integrin on their surface (**Figure S2F**). Consistent with the role of moesin in osteoclast fusion, moesin was the main activated ERM in these cells (**Figure 2F and S2C**). No significant difference was observed in the mRNA expression levels of osteoclast-marker genes in moesin KO compared with controls (**Figure S2G**), suggesting no alteration of osteoclast differentiation. Finally, the partial depletion of moesin by siRNA in human monocytes was also associated with a decline of P-ERM level (**Figure S2H-I**) and a significant increase in the fusion of hOCs (**Figure 2G-H**) in keeping with the findings obtained in mOCs. Together, these results imply that moesin restrains cell-cell fusion during the formation of multinucleated osteoclasts.

**Figure 2:**
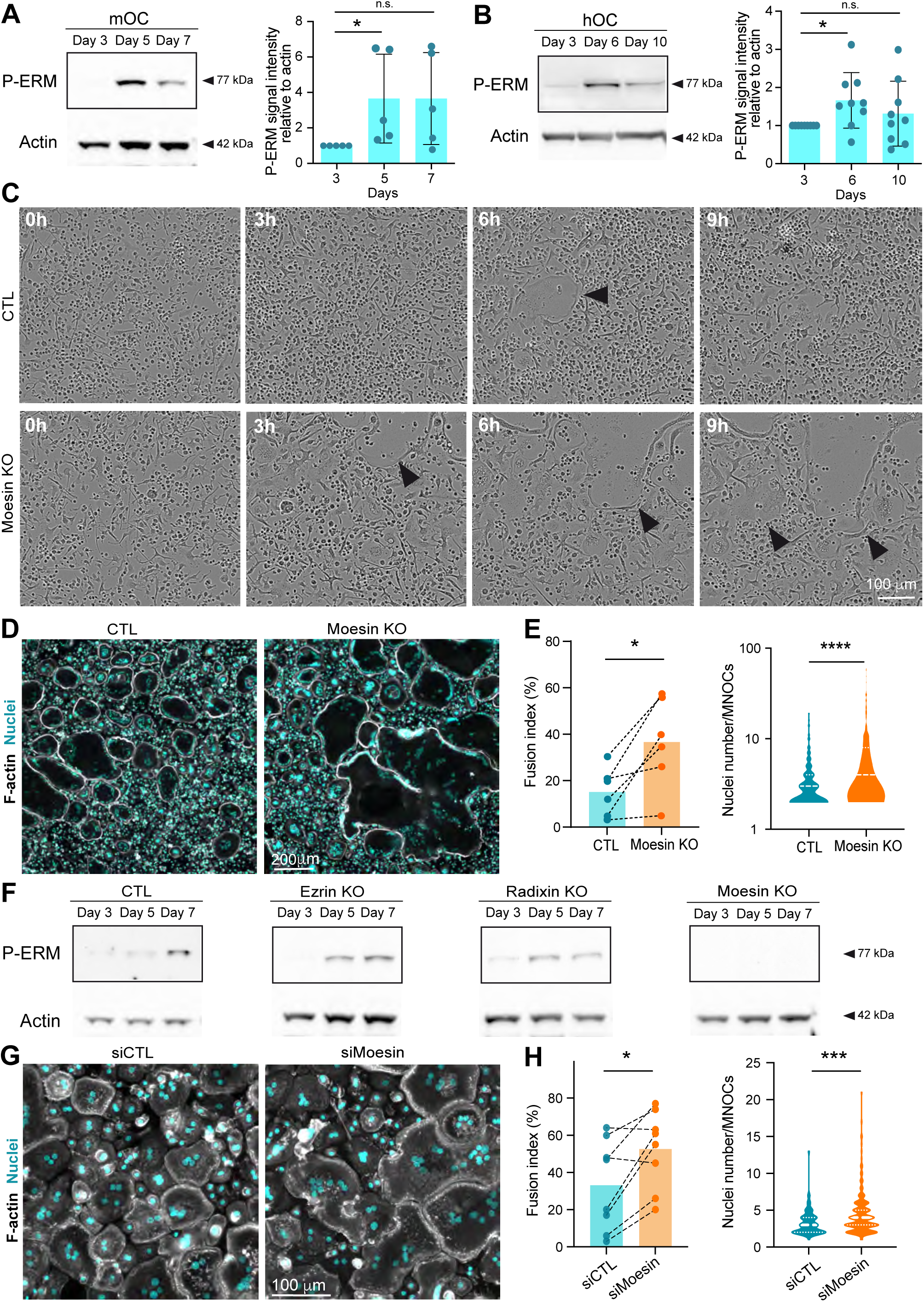
Moesin knock-out increases the fusion capacities of murine and human osteoclasts. **A.** Western blot analysis of activated ERM (P-ERM) expression level during mOC differentiation (day 3, 5 and 7); actin was used as loading control and quantification of P-ERM level was normalized to actin. Each circle represents an independent experiment, means ± SD are shown, n=5. **B.** Same experiment as in A during hOC differentiation (day 3, 6 and 10). Each circle represents a single donor, means ± SD are shown, n=9. (A-B) Predicted molecular weight are indicated. n.s. not significant. **C.** Reprensentative brightfield microscopy images from a time-lapse movie of control (CTL) and Moesin knock-out mOC (Moesin KO) at day 4 of differentiation. Black arrowheads point to multinucleated giant osteoclasts. See **Movie 6.** Scale bar, 100 µm. **D-E.** Microscopy analysis of cell fusion in control (CTL) and Moesin KO mOC. (D) Representative microscopy images: F-actin (phalloidin, white) and nuclei (DAPI, cyan). Scale bar, 200 µm. (E) Quantification of fusion index (n=6 independent experiments, 4 images/experiment, 1000 nuclei/image,) and nuclei number per multinucleated osteoclast (160-250 cells/condition, n=3). **F.** Representative Western blot analysis of P-ERM expression level in control (CTL), ezrin KO, radixin KO and moesin KO mOC; actin was used as loading control, n=2. Predicted molecular weight are indicated. **G-H.** Microscopy analysis of hOC fusion after treatment with non-targeting siRNA (siCTL) or siRNA targeting Moesin (siMoesin). (G) Representative microscopy images: F-actin (phalloidin, white) and nuclei (DAPI, cyan). Scale bar, 100 µm. (H) Quantification of fusion index (each circle represents a single donor, n=8) and nuclei number per multinucleated osteoclast (one representative experiment, 100-200 cells/condition, n=8).

To further investigate ERM activation in osteoclast fusion, we next synchronized this process using the hemifusion inhibitor lysophosphatidylcholine (LPC) that reversibly blocks membrane merging ^51–53^. Accumulation of ready-to-fused mononuclear cells correlated with an increase in the level of P-ERM (**Figure S3A-C,** +LPC). Following the washout of the drug, which synchronized fusion events, the phosphorylation of ERM returned to baseline levels (**Figure S3A-C**, +/-LPC), suggesting that reduced level of ERM activation promotes osteoclast fusion.

We next explored P-ERM levels in different pathological settings known to exacerbate osteoclast fusion such as during inflammation and upon HIV-1 infection ^54–56^. Murine osteoclasts derived from dendritic cells (DC-OCs) mimic osteoclasts in inflammatory conditions^57^, whereby they differentiate into osteoclasts containing more nuclei compared to those derived from monocytes (MN-OCs)^56^. P-ERM level was significantly diminished in DC-OCs compared to their “classical” osteoclast counterparts (**Figure S3D**) as well as in hOCs undergoing formation of HIV-1-positive giant syncytia (**Figure S3E-F**). Interestingly, the results were recapitulated in macrophages fusing upon HIV-1 infection ^58,59^ (**Figure S3G-H**), implying that the role of ERM activation in cell-cell fusion extends beyond osteoclasts. Together, these data indicate that the level of moesin activation is strongly correlated with the capacity of osteoclasts and macrophages to fuse in physiological and pathological contexts.

### Moesin depletion increases TNT formation and reduces membrane-cortex attachment

To explore the cellular mechanisms involved in the control of cell-cell fusion by moesin, we next monitored the subcellular localization of moesin and P-ERM (corresponding mainly to P-moesin) during osteoclast differentiation. In hOCs, moesin appeared associate to the plasma membrane, including at TNTs, zipper-like structures, podosome belts or sealing zones (**Figure S4-S5A** and **movie 7**). We also detected accumulation of P-ERM at the tips of a subset of TNTs (**Figure S5A**), leading us to characterize the impact of moesin depletion on the formation of TNTs. Interestingly, TNT formation was increased in the absence or after depletion of moesin in mOCs (**Figure 3A-B** and **S5B**) and hOCs (**Figure 3C-D** and **S5B**), respectively. In agreement with a specific role for thick TNTs (containing microtubules) in osteoclast fusion (**Figure 1**), we found that only the number of cells forming thick TNTs, and not thin TNTs, was affected by moesin depletion (**Figure 3B** and **3D**). Live imaging on 1:1 mixed cultures of Lifeact-Cherry-expressing control and Lifeact-GFP-expressing KO cells showed that cells form more TNT-like protrusions in the absence of moesin (**Figure S5C**).

**Figure 3:**
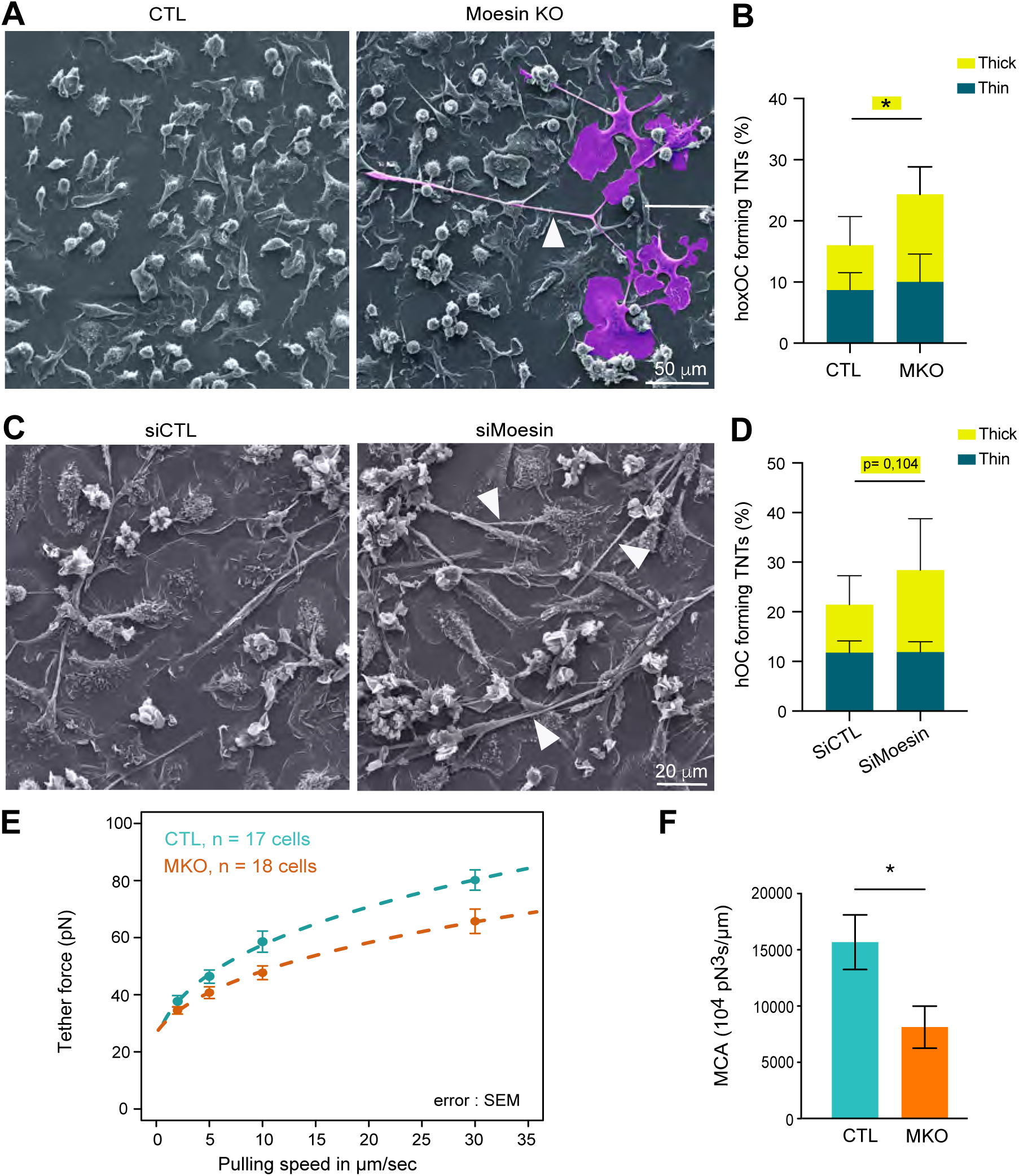
Moesin depletion enhances the formation of tunneling nanotubes (TNTs) and reduces membrane-cortex attachment. **A-D**. Effect of moesin depletion on TNT formation in mOCs (A-B) and hOCs (C-D). (A, C) Representative scanning electron microscopy images of mOCs (day 3) CTL *versus* Moesin KO (A) and hOCs (day 3) treated with non-targeting siRNA (siCTL) or targeting moesin (siMoesin) (C). White arrowheads show TNTs. (A) A giant mOC is colored in purple. Scale bar, 50 µm (A) and 20 µm (C). (B, D) Quantification of the percentage of cells forming thick and thin TNTs after immunofluorescence analysis in mOCs (B, n= 3 independent experiments) and hOCs (D, n=4 donors) (see Fig. S1B and methods), n>250 cells per conditions, means ± SD are shown. **E-F.** Analysis of force by atomic force spectroscopy operated in dynamic tether pulling mode. (E) Force-velocity curve from dynamic tether pulling on CTL and moesin KO (MKO) mOCs. Data points are mean tether force ± SEM at 2, 5, 10 and 30 µm/sec pulling velocity. At least 17 cells per condition were analyzed in 4 independent experiments. (F) Mean and standard deviation of the MCA parameter Alpha obtained from fitting the Brochard-Wyart model (see Methods for details).

ERM proteins regulate the physical properties of the membrane and the actomyosin cortex and control a plethora of cellular processes including the formation of cell protrusions ^60,61^. As such, we asked whether the physical link between the actomyosin cortex and the plasma membrane (Membrane-to-Cortex Attachment, MCA) was affected by the absence of moesin in mOCs, using atomic force microscopy-based force spectroscopy ^63^. Significantly lower forces were required to pull dynamic membrane tethers from moesin KO cells compared to controls (**Figure 3E-F**), corresponding to a 50% decrease in MCA after moesin depletion.

Thus, reduced levels of moesin is associated with a reduced attachment of the actin cytoskeleton to the plasma membrane, as well as an increased fusion and ability to form TNTs in osteoclasts. These data suggest that the depletion of the actin-membrane linker moesin promotes onset or stabilization of TNTs by decreasing MCA.

### Moesin depletion boosts bone degradation activity of osteoclasts

Because the bone degradative capacity of osteoclasts usually correlates with their multinucleation and size, we next examined the effect of moesin depletion on bone resorption. Accordingly, we found that mOCs differentiated from moesin KO precursors exhibited a ∼1.5-fold increase in bone resorption activity compared with control cells (**Figure 4A-B**). By performing mixed cultures of control-mcherry and KO moesin-GFP osteoclasts seeded on glass, we showed that the podosome belts (reminiscent of the sealing zones) were formed, for the majority, by moesin-depleted cells (**Figure S6A**). Of note, mixed color podosome belts were observed, consistent with the hypothesis that the fusion can occur between heterogeneous partners ^64,65^. We then assessed the number and the architecture of the sealing zone, which is crucial for bone resorption ^15,66^. In moesin KO mOCs seeded on bones, the total area covered by sealing zones was increased (**Figure 4C-D** upper panels, and **S6B**), corresponding to both an increase in the number and the surface covered by individual sealing zones, without any change in their circularity. In addition, the width of the F-actin-rich region inside the sealing zone was increased (**Figure 4C-D**, lower panels). As expected from the effect of moesin depletion on cell fusion, the number of nuclei inside cells forming sealing zones was increased upon moesin KO (**Figure S6C**). Consistently, siRNA-mediated silencing of moesin in hOCs recapitulated the effects of moesin KO on bone degradation and sealing zone formation (**Figure 4E-H**).

**Figure 4:**
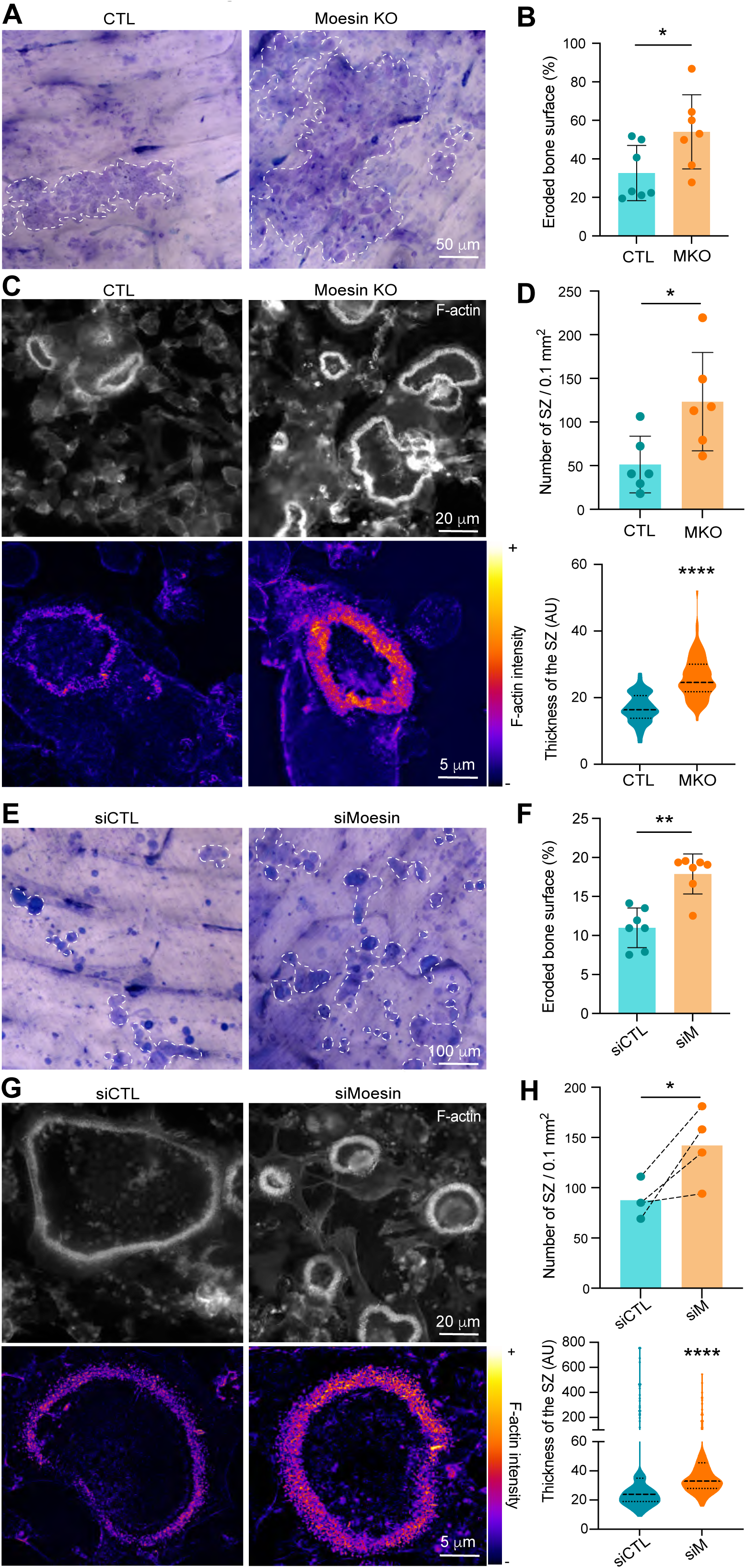
Moesin depletion boosts bone degradation in both murine and human osteoclasts. **A-D.** Effect of moesin KO on bone degradation (A-B) and sealing zone (SZ) formation (C-D) in mOCs. (A-B) mOC control (CTL) *versus* moesin KO (MKO) were cultured for 7 days on bone slices; after cell removal, bone was stained with toluidine blue. (A) Representative images of bone degradation, eroded bone surfaces are delineated by dashed white lines. Scale bar, 50 µm. (B) Quantification of bone eroded surface (%) using semi-automatic quantification (n=3 independent experiments, 2 or 3/bone slices/condition, means ± SD are shown). (C-D) mOC control (CTL) versus moesin KO (MKO) were cultured for 5 days on glass coverslips, detached and then seeded for additional 2 days on bone slices. (C) Representative microscopy images of sealing zones visualized by F-actin staining (phalloidin, white in upper panels and colored-coded intensity in lower panels). Scale bars, 20 and 5 µm. (D) Quantification of the number of sealing zones per bone surface (n=3 independent experiments, 2 bone slices/condition, means ± SD are shown) and of sealing zone thickness (n=3 independent experiments, 15-20 SZ/condition, 3 locations/SZ). **E-H.** Effect of moesin depletion on bone degradation (E-F) and sealing zone formation (G-H) in hOCs. 6 day-differentiated hOCs on glass coverslips treated at day 0 with siCTL or siMoesin (siM) were detached and seeded for additional 24h on bone slices. (E) Same legend as in A. Scale bar, 100 µm. (F) Same legend as in B (n=4 donors, 2 bone slices/condition, SD are shown). (G) Same legend as in C. (H) Same legend as in D. Quantification of the number of sealing zones (n=4 donors, 2 bone slices/condition) and of sealing zone thickness (n=3 donors, 15-20 SZ/condition, 3 locations/SZ).

To investigate whether moesin regulates bone degradation in addition to its role in osteoclast fusion, we uncoupled these two processes. To do so, we depleted moesin using siRNA in already multinucleated mature hOCs (**Figure S7A**) and found no effect on the fusion index (**Figure S7B**). However, under these conditions, the level of expression of moesin and of P-ERM was reduced (**Figure S7C**), which coincided with an increase in bone degradation (**Figure S7D)**. Of note, the two main modes of bone resorption (i.e. pits and trenches) made by hOCs^67^ were not differentially affected (**Figure S7E**). Finally, depletion of moesin in late stage of hOC differentiation also favored the formation of sealing zones and increased their thickness (**Figure S7G-H**). Thus, moesin inhibits osteoclast activity at two levels, (i) by controlling the fusion capacity of osteoclasts and (ii) by regulating sealing zone number and structure modulating the efficiency of the bone degradation machinery.

### The RhoA/SLK axis acts downstream of β3-integrin to control ERM activation in osteoclasts

Next, we explored by which signaling pathway moesin activation regulates the formation of the sealing zone in mature osteoclasts. Key regulators of actin dynamics known to regulate podosome and sealing zone dynamics include the small GTPases of the Rho family ^68,69^, RhoA, Rac1/2 and Cdc42 ^14,70–72^. RhoGTPase-dependent signaling pathways are also known to regulate the ERM protein activation cycle in other cell types ^29,73,74^. First, we tested whether pharmacological inhibition of RhoGTPases affect the activation status of ERM proteins in hOCs. For this, we used the exoenzyme C3 transferase (TATC3), NSC23766, and ML141 that target RhoA, Rac1/2 and Cdc42, respectively. Compared to the strong effects of calyculin A and staurosporine, used as positive and negative controls, respectively (**Figure S8A**), we observed a significant decrease in P-ERM levels only after TATC3 treatment (**Figure 5A and S8B**), suggesting that RhoA is the main small GTPase involved in ERM activation in osteoclasts. Two Ser/Thr kinases, SLK and ROCK, have been described to be activated by RhoA ^75–77^ and to directly phosphorylate ERM proteins ^78–80^. Treatment with Y-27632, which inhibits ROCK1 and ROCK2, did not impact the level of P-ERM (**Figure S8C**) while downregulation of SLK by siRNA resulted in a significant decrease in ERM activation (**Figure 5B**). Accordingly, the sealing zones in SLK-deficient osteoclasts are thicker than the controls (**Figure 5E-F**), mimicking the effect of moesin depletion (see **Figure 4**).

**Figure 5:**
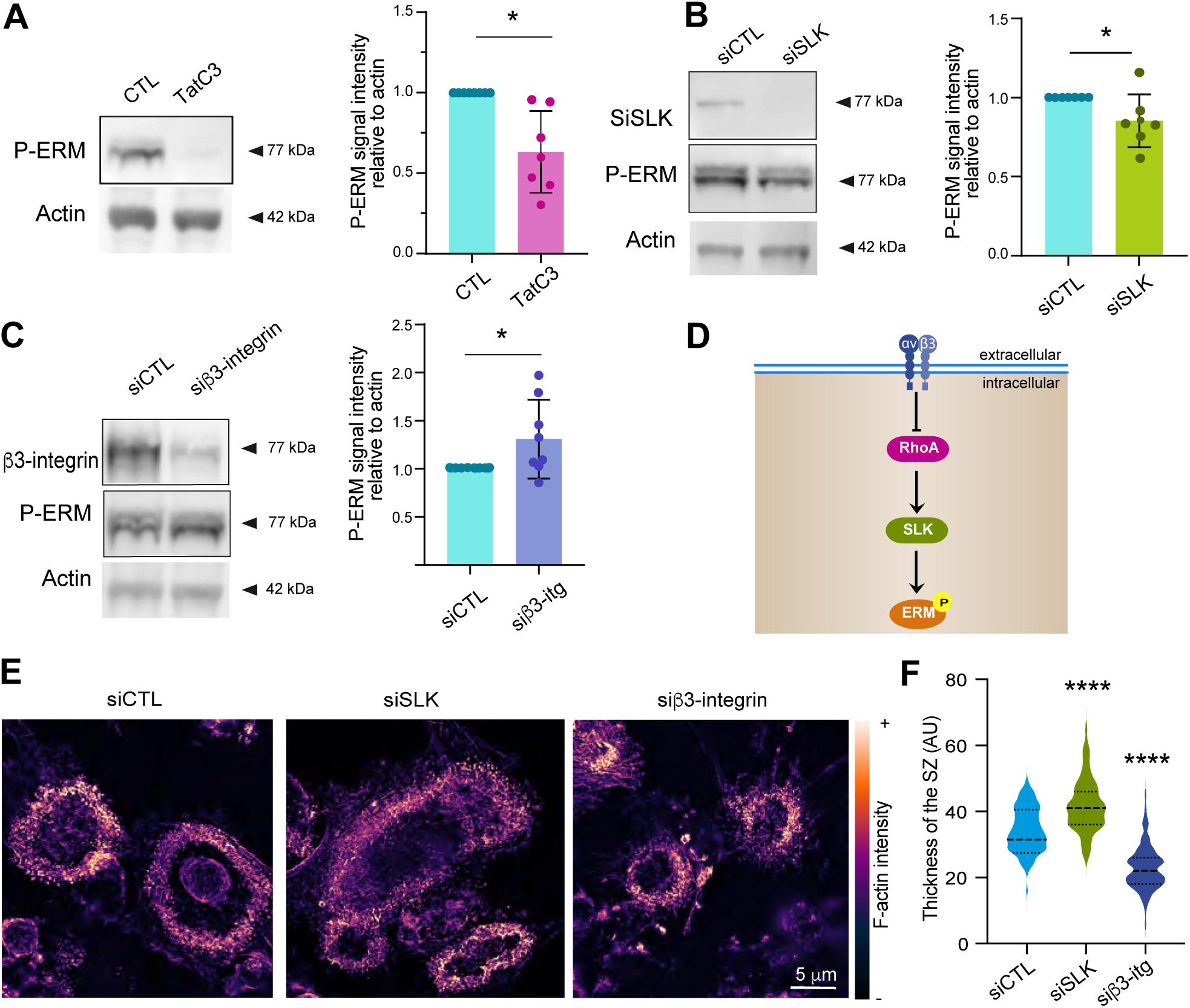
The Rho/SLK axis downstream of β3-integrin controls ERM activation and sealing zone formation. **A.** RhoA inhibition reduces ERM activation. 6 day-hOCs were treated or not (CTL) with TatC3, targeting the RhoGTPases RhoA. Representative Western blot analysis (left) and quantification of P-ERM signal normalized to actin (right). Each circle represents a single donor (n=6, means ± SD are shown). **B.** SLK suppression reduces ERM activation. hOCs were treated with non-targeting siRNA (siCTL) or siRNA targeting SLK kinase (siSLK). Representative Western blot analysis (left) and quantification of P-ERM signal normalized to actin (right). Each circle represents a single donor (n=7, means ± SD are shown). **C.** β3-integrin suppression favors ERM activation. hOCs were treated with non-targeting siRNA (siCTL) or siRNA targeting β3-integrin (siβ3-integrin). Representative Western blot analysis (left) and quantification of P-ERM signal normalized to actin (right). Each circle represents a single donor (n=7, SD are shown). (A-C) Predicted molecular weight are indicated. **D.** Schematics showing the proposed Rho/SLK axis downstream of β3-integrin for ERM activation**. E-F.** Effect of SLK and β3-integrin depletion on the formation of sealing zones in hOCs. (E) Representative super-resolution microscopy images of sealing zones visualized by colored-coded intensity of F-actin (phalloidin, Scale bar, 5 µm) and (F) quantification of sealing zone thickness (n=2 donors, 15-20 cells/condition and 3 locations/SZ).

Finally, to determine the signal that could trigger RhoA/SLK-dependent ERM regulation, we tested the importance of αvβ3 integrin. Indeed, this marker of mature osteoclasts ^81,82^ mediates their ability to polarize, spread and degrade bone ^70,83–85^. We showed that β3-integrin depletion (**Figure 5C**) affects the formation of the sealing zones confirming previous observations ^70^, with a decrease in their width (**Figure 5E-F**). Importantly, phosphorylation of ERM proteins was enhanced in β3-integrin-depleted cells (**Figure 5C**). Altogether, these results provide evidence that, in mature osteoclasts, ERM activation and sealing zone formation is under the control of the RhoA/SLK axis, downstream of the β3-integrin (**Figure 5D**).

### Mice lacking moesin exhibit bone loss and increased osteoclast number and activity

Finally, to explore the physiological relevance of moesin to osteoclast and bone biology, we assessed moesin expression and function in long bones of mice. As shown by immunohistology experiments on serial sections of femur of wild type (WT) mice, moesin is expressed in cathepsin K-positive osteoclasts lining bone surface (**Figure S9A**), in addition to other cells residing within the bone marrow. We next examined the bone phenotype of moesin global knockout mice (*Msn*-/-) ^35^. No difference in the size, weight or skeleton of matched littermates up to 40-weeks-of-age was observed (**Figure S9B**). Nonetheless, micro-computed tomographic (μCT) analysis of the distal femurs of 10-week-old male WT and *Msn*-/- mice revealed that the long bones of null mice exhibited trabecular bone loss (**Figure 6A**), as quantified by a significantly lower trabecular bone surface fraction (BS/TV), trabecular thickness and trabecular number, associated with an increase in trabecular separation, compared to WT mice (**Figure 6B**). Thus, moesin-deficient mice are osteopenic. In these mice, cortical bone parameters were not affected (**Figure S9C**). Importantly, histological analysis showed that the TRAP-positive signal of osteoclasts was significantly increased in bones of *Msn*-/- mice compared to WT (**Figure 6C-D**), demonstrating that the deletion of moesin increases osteoclast number and activity in bones.

**Figure 6:**
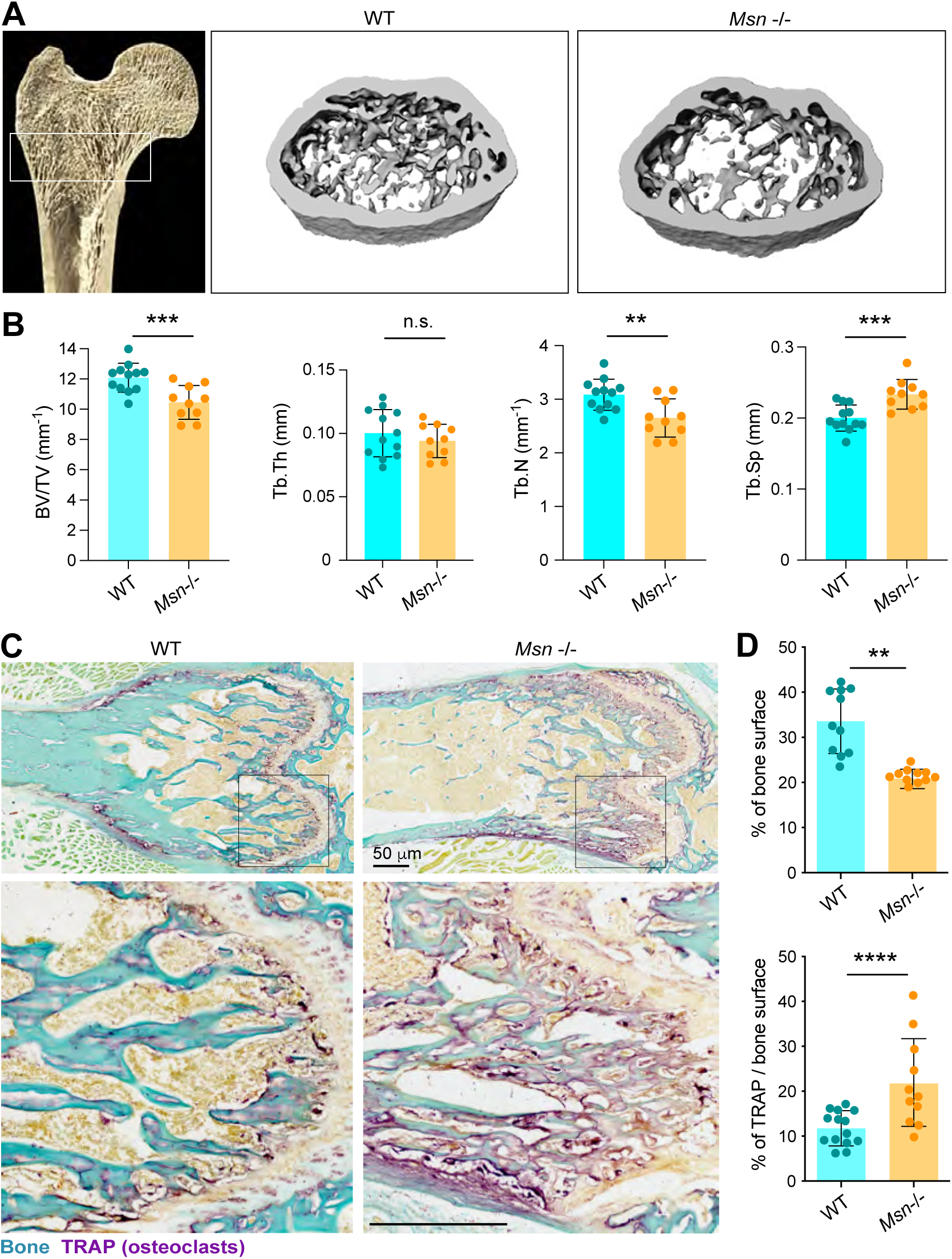
Moesin deficiency translates in bone defects and high osteoclast activity in vivo. **A.** Representative microcomputed tomography images of trabecular section of distal femurs from WT mice and *Msn*^-/-^ mice. **B.** Histograms indicate means ± SD of trabecular bone volume per total volume (BV/TV), trabecular thickness (Tb.Th), number (Tb.N) and separation(Tb.Sp). Animal groups were composed of 6 mice of each genotype, and 11 femora were analyzed in total for *Msn*^-/-^ mice and 9 femora for the WT mice group. n.s. not significant. **C.** Histological analysis of osteoclasts using TRAP staining (osteoclast in purple) and fast green (bone is green) on femurs from WT mice and Msn-/- mice. Scale bars: 50 µm. **D.** Histogram indicates the mean ± SD of bone surface (upper panel) and TRAP+ surface per bone surface (lower panel) for each condition. N ≥ 2-3 sections chosen among the most median part of 4 mice for each genotype. Each circle represents a single bone section.

## DISCUSSION

Here, we show that ERM activation, specifically moesin activation, plays a negative regulatory role in osteoclast formation and bone resorption. First, we demonstrate that it acts during the early stages of osteoclastogenesis, by regulating cell-cell fusion via the formation and/or stabilization of TNTs. Second, in mature osteoclasts, activation of moesin, under the control of the β3-integrin/RhoA/SLK axis, regulates osteolysis by impacting the number and structure of the sealing zones. Related to osteoclast dysfunction *in vitro*, mice bearing total *moesin* deletion develop an osteopenic phenotype with increased osteoclast activity. This phenotype is observed in trabecular bones but not cortical ones that are less remodeled in adult mice. Given the intricate interplay between osteoclasts and other bone cells and the ubiquitous expression of moesin, we cannot exclude that moesin also exerts regulatory effects in other cells. However, the preferential effect of moesin on osteoclast activity over differentiation may make it a relevant candidate to control bone loss.

We first demonstrate that moesin activation acts as a novel regulatory mechanism that limits the extent of osteoclast fusion, preventing an excessive number of nuclei per osteoclast, and thus insuring optimal osteolytic activity. Consistent with this hypothesis, we demonstrate that the level of osteoclast fusion under pathological or drug-induced conditions is negatively correlated with the level of ERM activation; LPC-dependent fusion inhibition is associated with enhanced ERM activation while fusion increase in inflammatory context or upon HIV-1 infection is associated with decreased ERM phosphorylation. Moreover, macrophage fusion induced by HIV-1 infection also correlates with downregulation of P-ERM levels, suggesting that the inhibitory effect of ERM activation during cell-cell fusion extends to other myeloid cell types. It would be interesting to know whether phosphorylation of ERM proteins serves as a general regulator of membrane fusion, for example in the formation of myofibers or syncytiotrophoblasts that do not necessarily involve TNT-like structures ^3,86^. Depending on the cell type, ERMs have been identified as inhibitors or boosters of the cell fusion process, e.g. activated ezrin prevents the formation of HIV-1 induced T cell syncytia while it promotes trophoblast and myotube fusion ^39,40,87–89^. Here, we propose that moesin activation is a novel regulatory mechanism that limits the extent of osteoclast fusion, preventing an excessive number of nuclei per osteoclast thus insuring optimal osteolytic activity.

We then propose that moesin controls the fusion of osteoclasts by limiting the number of thick TNTs. First, our live cell imaging studies provide definitive proof that TNTs play a critical role in osteoclast fusion, as previously suggested ^3,20–22,90^. Consistently, a peak in the number of cells emitting TNTs precedes the fusion process. Second, our data strongly suggest that moesin activation controls TNT number. This novel function of moesin is consistent with the well-known role of ERM proteins in the formation of actin-rich protrusions such as filopodia and microvilli ^60,91–93^. Indeed, P-ERM is localized over the entire surface of TNTs where we found cell fusion to occur. How TNT formation contributes to the fusion process remains speculative. A simple explanation could be that the more TNTs are present, the more membrane surface is available for fusion events to occur. Another tempting hypothesis is that TNTs serve as membrane platforms to bridge connections between distal cells, not necessarily the closest, but potentially between ideal fusion-competent partners ^94^. We also highlight the exclusive involvement of thick TNTs, which allow the microtubule-dependent transport of material between two connected cells ^24^, in moesin-dependent osteoclast fusion. Thick TNT formation decreases during osteoclast maturation and increases in moesin-depleted cells, showing a correlation between the fusion extent and the number of this subtype of TNTs. This is coherent with intercellular transports of molecules essential to the fusion process, such as phospholipids and DC-STAMP, which occurs through TNTs in osteoclast precursors ^20^.

Consistent with the role of ERM proteins in connecting the plasma membrane to the cortical actin network in many cell types ^95,96^, depletion of moesin in osteoclast precursors strongly decreases membrane-to-cortex attachment while the number of TNT-forming cells and cell-cell fusion increase. Thus, the release of actin from the membrane seems to be a favorable condition for the fusion process, either indirectly by supporting TNT onset, or directly by contributing to membrane fusion. As an example of a direct contribution, low membrane tension induced by myosin IIA reduction allows osteoclast fusion ^97^. From our results, we propose that during the early steps of osteoclastogenesis, a low ERM activity promotes the formation of actin-protrusions favorable to cell-cell fusion, then, and as soon as the proper number of nuclei per cell is reached, moesin is activated and counteracts the fusion process.

We show that moesin acts at a second step during osteoclastogenesis. In addition to regulating osteoclast fusion, moesin also influences the number and architecture of the sealing zones. These structures are composed of a dense network of podosomes organized in clusters ^14,16^ and as defined in macrophages, each individual podosome can exert a protrusion force on the substrate that is correlated to the F-actin content ^98^. Thus, the increased width of the podosome-rich zone observed upon moesin depletion may favor an efficient sealing of osteoclasts to the bone and therefore increase the concentration of bone-degradative molecules in the resorption area ^14,82^. Consequently, the exacerbation of bone resorption observed after moesin deficiency could result not only from the increase in the number and surface area of the sealing zones, but also from the ability of osteoclasts to adhere to the bone. Phosphorylation of moesin have been shown to be important for podosome rosette formation in Src-transformed fibroblasts ^99^ while in pancreatic cancer cells, ezrin regulates podosome organization independently of its activation ^100^. As mentioned previously, ERM proteins crosslink the actin cytoskeleton to several transmembrane proteins including CD44 ^101,102^, which might participate in the organization of the sealing zone. Indeed, in addition to its role in cell fusion ^3^, CD44 localizes to podosome cores and participates in podosome belt patterning in osteoclasts ^103^.

In mature osteoclasts, we showed that ERM activation depends on the RhoA/SLK axis. Such a mechanism of ERM regulation has already been described in several contexts, including the cell rounding at mitotic entry of dividing cells or the formation of the apical domain of epithelial cells ^29,93^. Moreover, we identified β3-integrin as an upstream regulator of this pathway. This marker of mature osteoclasts ^81,82^ mediates their ability to polarize, spread and degrade bone ^70,83–85^. We propose that β3-integrin limits the phosphorylation of moesin through the inhibition of the RhoA/SLK axis and in this way controls the number/architecture of the sealing zones. Moesin could be a new effector of the β3-integrin /Rho pathway, acting as a complementary regulatory mechanism to those already described ^70,85^.

In conclusion, in addition to the well-characterized role of ERM proteins in cell polarization and migration, this study provides evidence for a new role of moesin in osteoclast formation and function, including in vivo, by controlling fusion events and osteolytic activity. In osteoclasts, moesin is a key actin-structure regulator, regulating both actin-protrusive TNTs and podosome organization in the sealing zones. Targeting this protein or its regulatory pathway may present an opportunity to modulate the activity of osteoclasts without affecting their viability or differentiation, and thus may represent a potential target for the treatment of osteoclast-related bone diseases.

## METHODS

Materials and methods, chemical and antibodies, cell culture and transfection, HIV-1 infection, immunoblotting, qPCR, immunofluorescence, live imaging and microscopy, including scanning and atomic force microscopy, flow cytometry, bone resorption assays, histological and µCT analyses of mice bones, and statistical analysis are described in supplemental information, supplemental Methods.

Human monocytes were provided by Etablissement Français du Sang, Toulouse, France, under contract 21/PLER/TOU/IPBS01/2013–0042. *Msn* -/- mice were housed under barrier conditions in the Children’s Hospital of Philadelphia animal facility, in accordance with protocols approved by the Institutional Animal Care and Use Committee.

## ACKNOWLEDGEMENTS

We acknowledge the TRI imaging facility, member of the national infrastructure France-BioImaging supported by the French National Research Agency (ANR-10-INBS-04), in particular Isabelle Fourqueaux (CMEAB), Emmanuelle Näser, Eve Pitot and Elodie Vega (IPBS). We thank the multi-pathogen BSL3 laboratory. We thank the Wellcome Sanger Institute for providing a mouse line expressing Cas9 nuclease. We also thank Anne Blangy and Jean-Luc Davignon for fruitful discussions. This work was supported by the *Centre National de la Recherche Scientifique*, *Université Toulouse III - Paul Sabatier (UT3)*, the *Institut National de la Santé et de la Recherche Médicale*, the *Agence Nationale de la Recherche* (ANR16-CE13-0005-01, ANR DFG 2020 JA-3038/2-1 and ANR-20-CE14-0037)), the *Fondation pour la Recherche Médicale* (DEQ2016 0334894 ; DEQ2016 0334902, ENV202003011510), the *Fondation Toulouse Cancer Santé* and l’INSERM Plan Cancer, and the Penn Center for Musculoskeletal Disorders (NIH/NIAMS P30 AR069619). This study has been partially supported through the grant EUR CARe N°ANR-18-EURE-0003 in the framework of the Programme des Investissements d’Avenir. This work was also supported by NIH R01 AI147118 to JKB, CIHR (PJT-1620109) to SC, the European Molecular Biology Laboratory (EMBL) to AD-M, and the ATIP program (CNRS INSB) and the *Fondation pour la recherche contre le cancer* to FL. NJP is supported by funding from the *National Health & Medical Research Council of Australia* (APP2029078 and APP2020097). OD was supported by doctoral scholarships from *Université Paul Sabatier*, *Fondation pour la recherche contre le cancer (ARC*) and the Foundation F. Initiativas for the Minerva Trophy. PV, TS and MP were supported by doctoral scholarships from *Université Toulouse III - Paul Sabatier (UT3), La ligue contre le cancer* with *Fondation pour la recherche contre le cancer*, and the EUR CARe graduate school, respectively. CJP is supported by NIH/NIGMS 2K12GM081259-16.

## AUTHORSHIP CONTRIBUTION

OD performed all the experiments, with the help of PV, MP, TS, JH, RM, RG, GA and MBN. OD, BRM and CV designed the experiments and analyzed the results. MB participated in MCA experiments. PV generated the KO cells. MJ collected the bones. AM performed histological analysis. CJP and JDB assisted with microcomputed tomography experiments. BRM and CV supervised the project. RP, FL, VLC, SB and JKB co-supervised the project. IMP, NJP, ADM, FL, SC, CBW, RP and JKB obtained funding. BRM and CV wrote the manuscript with input from the other authors.

