## Supplemental material for "Moesin controls cell-cell fusion and osteoclast function"

**SUPPLEMENTAL INFORMATION**

#### SUPPLEMENTAL METHODS

***Mice.*** *Moesin* (-/-) (*Msn-/-*) mice, backcrossed onto the C57Bl6/J background, were previously characterized [1]. Briefly, KO mice were generated by and purchased from the Texas A&M Institute for Genomic Medicine. A gene trap vector was inserted into the first intron of the *Msn* gene on the X chromosome in 129/Sv ES clones (OST432827), and live mice with germline insertion were generated on the 129Sv × C57BL/6 background. The resulting KO mice were backcrossed for 10 generations to mice of the C57BL/6 background (The Jackson Laboratory). All mice were housed under barrier conditions in the Children's Hospital of Philadelphia animal facility, in accordance with protocols approved by the Institutional Animal Care and Use Committee.

***Chemicals and antibodies.*** Human recombinant macrophage colony-stimulating factor (M-CSF) was purchased from Peprotech (London, UK) and human receptor activator of NF-κB-ligand (RANKL) and mouse M-CSF and RANKL (mouse and human) were from Miltenyi Biotec (Germany). DAPI was purchased from Sigma-Aldrich (Lyon, France). The following rabbit antibodies were from Cell Signaling: anti-integrin β3 (#4702), anti-ezrin (#3145), anti-radixin (#2636), anti-moesin (#3150), rabbit anti-phospho-ERM (#3141). α-tubulin (T9026, clone DMA1, Sigma-Aldrich), anti-HIV p24 (KC57-FITC, clone FH190-1-1, mouse IgG1, Beckman Coulter, 6604665), rabbit anti-actin (A5060, Sigma-Aldrich) and anti-vinculin (V9131, clone HVIN-1, Sigma Aldrich) were also used. Secondary HRP-conjugated antibodies were from Dako and fluorescent secondary antibodies and Texas Red/Alexa Fluor 488-conjugated phalloidins were obtained from Molecular probes (Invitrogen, Cergy Pontoise, France).

Inhibitors of the signaling pathway Rho/ROCK were Y27632 (50 µM, Sigma), C3 exoenzyme coupled to permeant peptide TAT (TAT-C3, 10 µg/mL), produced in G. Fabre laboratory as previously described (Sahai and Olson, 2006), NSC23766 (100µM, ab142161, Abcam) and ML141 (10nM, SML0407, Sigma Aldrich). The phosphatase inhibitor calyculin A (Sigma) and the protein kinase C inhibitor staurosporine (Sigma) were used at 100nM. TAT-C3 was added at 48h, Y27632 at 24h, NSC23766 and ML141 at 10h, and calyculin and staurosporine at 10 min. Osteoclast fusion synchronization with LPC (1-lauroyl-2-hydroxy-sn-glycero-3-phosphocholine, Avanti Polar Lipids #855475) was performed as described in[2]. Briefly, at day 6 of differentiation media was refreshed with medium containing M-CSF, RANKL and 350 µM LPC. Following 17 h treatment, LPC was washed and cells were maintained in a fresh media for additional 90 min.

***Bone histomorphometric analysis.*** Bones from 10-week-old wild-type (WT) and *Msn-/-* (*Msn*$^{-/)y}$ male littermate mice were fixed in PBS plus 4% paraformaldehyde overnight at 4°C and then washed and stored in 70% ethanol. Bone microarchitecture analysis using high-resolution µCT was performed at the pre-clinical platform ECELLFRANCE (IRMB, Montpellier, France). Cortical and trabecular femora were imaged using high-resolution µCT with a fixed isotropic voxel size of 9µm with X-ray energy of 50 kV, current of 500 mA, 0.5 mm aluminum filter, and 210ms exposure time. Quantification of bone parameters was performed on the trabecular region of the proximal part of each femur (172 mm long) and on the cortical region (0.43 mm long region centered at the femoral midshaft) on CT Analyzer software (Bruker microCT, Belgium). For visual representation, 3D reconstructions (8.8-mm cubic resolution), were generated using NRecon software (Bruker µCT, Belgium). Animal groups were composed of 6 mice each, and 11 femora were analyzed in total for *Msn-/-* mice and 9 femora for the WT mouse group.

Whole skeletons were analyzed using the Scanco vivaCT80 (Nokomis, FL, USA). Immediately after euthanasia, animals were scanned using a voxel size of 100µm, X-ray energy of 55 kVp, current of 145 µA and integration time of 300 ms with a 0.5 mm Al filter.

***Histological analysis.*** For immunohistofluorescence of frozen bone sections (10 µm, Leica CM1950), femurs from B6 mice were fixed with 4% paraformaldehyde (Electron microscopy science 157-4) overnight at 4°C, decalcified 10 days with 10% EDTA (Sigma ED4SS) changed daily, and then incubated in 30% sucrose (Euromedex 200-301-B) solution overnight at 4°C prior to embedding in OCT (CellPath KMA-0100 00A) and snap-freezing in isopentane (Sigma M32631) pre-cooled by liquid nitrogen. After saturation and permeabilization/blocking for 1 hour at room temperature with 5% goat serum (Genetex GTX73249), 5% BSA (Euromedex 04-100-812-C), 0,2% Triton X-100 (Sigma T8532), the sections were stained overnight at 4°C with antibodies to moesin (Q480, rabbit, 1:100; Cell Signaling 3150) or anti-cathepsin K (1:100; Abcam ab19027). Goat anti-rabbit Alexa Fluor 555 (1:400; Cell Signaling #4413) secondary antibodies was used. Nuclei were visualized with 49,6-diaminidino-2- phenylindole (DAPI; Sigma D9542). Images were acquired using a Zeiss Axio Imager M2 using an X40/0.95 Plan Apochromat objective (Zeiss) and an ORCA-flash 4.0 LT (Hamamatsu) camera, and processed using the Zeiss Zen software.

For TRAP staining, femurs and tibia from adult WT and *Msn-/-* mice male littermate mice were fixed in PBS plus 4% paraformaldehyde overnight at 4°C, decalcified in EDTA, and freezed in OCT (CellPath KMA-0100-00A). Longitudinal serial 10µm cryosections of the median portion of whole bone were stained for TRAP (Sigma 386A) and by fast green (Sigma F7252). Images were acquired using a Zeiss Axio Imager M2 using an X10/0.3 Plan neofluar objective (Zeiss) and an AxioCam 503 color (Zeiss) camera, and processed using the Zeiss Zen software. The percentage of TRAP-positive staining by bone surface and the percentage of bone surface by surface of ROI analysed were quantified with QuPtah software. The stainings were quantified on ≥ 2-3 sections chosen among the most median part of 4 mice for each genotype.

***Human osteoclasts (hOCs) and RNA interference.*** Human peripheral blood mononuclear cells were isolated from the blood of healthy donor buffy coats (Etablissement Français du Sang, Toulouse, France, contract 28 21/PLER/TOU/IPBS01/20130042), as described previously [3]. Briefly, cells were centrifuged through Ficoll-Paque Plus (Dutscher), resuspended in cold phosphate buffered saline (PBS) supplemented with 2 mM EDTA, 0.5% heat-inactivated Fetal Calf Serum (FCS) at pH 7.4 and monocytes were sorted with magnetic microbeads coupled with antibodies directed against CD14 (Miltenyi Biotec # 130-050-201). For differentiation to hOCs, monocytes were seeded on slides in 24-well plates at a density of $5x10^5$ cells per well in RPMI supplemented with 10% FCS, human M-CSF (50 ng/mL) and human RANKL (30 ng/mL). The medium was replaced every 3 days with medium containing h-M-CSF (25 ng/mL) and h-RANKL (100 ng/mL). hOCs from the same donor were used from day 1 to 3 of differentiation (osteoclast precursors) or at day 6 to 10 of differentiation (mature osteoclasts). CD14+ human monocytes were transfected with 200 nM siRNA using the HiPerfect system (Qiagen) as described previously. The mix of HiPerfect and siRNA were incubated for 15 min at room temperature and then the cells were added drop by drop. To deplete Moesin in the late stage of osteoclast differentiation, siRNA transfection was performed at day 6. The following siRNA (Dharmacon) were used: human ON-TARGET plus SMART pool siRNA non-targeting control pool (siCTL); human ON-TARGET plus SMART pool siRNA targeting MSN (Moesin) sequences: 5'-CGUAUGCUGUCCAGUCUAA-3'; 5'-GAGGGAAGUUUGGUUCUUU-3'; 5'-UCGCAAGCCUGAUACCAUU-3'; 5'-GGCUGAAACUCAAUAAGAA-3'. The human ON-TARGET plus SMART pool siRNA targeting ITGB3 (β3-integrin) sequences :5'-GCFUGAAUUGUACCUAUA-3' ; 5'-GAAGAACGCGCCAGAGCAA-3' . 5'-GCCAACAACCCACUGUAUA-3' ; 5'-CCAGAUGCCUGCACCUUUA-3'. The human ON-TARGET plus SMART pool siRNA targeting SLK sequences: 5'-GGUAGAGAUUGACAUAUUA-3' ; 5'-GAAAAGAGCUCAUGAAACG -3'; 5'-GCUCGAAGAACGACACUUA -3'; 5'-GGAACAUAGCCAAGAAUUA-3'

***HIV-1 infection of hOCs and macrophages***. At day 6 of differentiation, hOCs or human monocyte-derived macrophages were infected with the viral strain NLAD8-VSVG, produced by cotransfection with the proviral plasmid in combination with pVSVG (from S. Bénichou laboratory), as described in [4]. Cells were harvested 7 days post-infection.

***Murine HoxB8-derived osteoclasts (mOCs) and generation of single Knockout for each ERM protein. .*** Myeloid progenitors were isolated from the bone marrow of a mouse carrying the EF1a-hCas9-IRES-neo transgene in the ROSA26 locus [5], and immortalized by transduction with a retrovirus allowing conditional expression of the HoxB8 homeobox gene, as previously ddescribed [6]. HoxB8-derived progenitor cells were cultured in complete medium consisting of RPMI-1640 medium (GIBCO) with 10% Fetal Bovine Serum (FBS, Sigma-Aldrich), 2 mM L-glutamine, 100 U/mL penicillin, and 100 µg/mL streptomycin (GIBCO) and supplemented with 20 ng/mL GM-CSF (Miltenyi Biotec) and 5 µM β-Estradiol (Sigma-Aldrich), as. For differentiation in osteoclasts (called mOCs, for mouse osteoclasts), HoxB8 progenitors were collected, washed twice in complete medium and seeded on glass coverslips (Paul Marienfeld GmbH) in 12-well plates (1,8x$10^5$cell/well for CTL, KO Ezrin and KO Radixin cells ; 1,2x $10^5$ for KO Moesin cells) osteoclasts were differentiated in complete RPMI medium supplemented with 25 ng/mL murine M-CSF and 100 ng/mL murine RANKL (Miltenyi Biotech). The cells were incubated at 37°C in a 5% CO2 incubator. Medium and cytokines were replaced every 3 days and mature osteoclasts were obtained at day 5-7.

For the generation of single knockout cell lines for each ERM protein, the sgRNAs used were for EZR: (5'-CTACCCCGAAGACGTGGCCG-3'), RDX: (5'-GCCATCCAGCCCAATACAAC-3'), and MSN: (5'- TATGCCGTCCAGTCTAAGTA-3'). We targeted luciferase as a control, with the sgRNA LUC: (5'- GGCGCGGTCGGTAAAGTTGT-3'). The sgRNAs were introduced into the pLenti-sgRNA backbone (Addgene #71409) with digestion by BsmBI (New Englands Biolabs #R0739) and T4 DNA ligase (Fischer Scientific EL011L). For lentiviral particle production, Lenti-X 293T cells (Clontech) were co-transfected with PMDL (Addgene #12251), REV (Addgene #12253), VSVG (Addgene #12259) and specific sgRNA-cloned plasmids using Lipofectamine 3000 and OptiMEM according to the manufacturer's guidelines [7]. Then, 2×$10^5$ Cas9-expressing HoxB8 progenitors were transduced with viral particles and Lentiblast Premium (OZ Biosciences). After 24h, the transduced cells were selected with 10 µg/mL of puromycin (Invivogen) for 2 days and validated by immunoblotting.

***Bone marrow dendritic-cell derived osteoclasts.*** DC-derived (DC-OCs) and monocyte-derived (MN-OCs) osteoclasts were differentiated in vitro as described previously [8–10] from 6-week-old C57BL/6 mice. Briefly, CD11c$^+$ BM-derived DCs were obtained by culturing 5×$10^5$ BM cells/well in 24-well plates in RPMI medium (ThermoFisher Scientific) supplemented with 5% serum (Hyclone, GE Healthcare), 1% penicillin-streptomycin (ThermoFisher Scientific), 50 µM 2-mercaptoethanol (ThermoFisher Scientific), 10 ng/mL GM-CSF, and 10 ng/mL IL-4 (both from PeproTech). CD11c+ DCs were isolated using biotinylated anti-CD11c (1:200; clone HL3; BD Biosciences) and anti-biotin microbeads (Miltenyi Biotec). DC-OCs were differentiated by seeding a total of 2×$10^4$ CD11c$^+$ DCs/well on 24-well plates in MEM-alpha (ThermoFisher Scientific) including 5% serum (Hyclone, GE Healthcare), 1% penicillin-streptomycin, 50 µM 2-mercaptoethanol, 25 ng/mL M-CSF, and 30 ng/mL RANK-L (both from R&D) (OC differentiation medium). For MN-OCs culture, 2×$10^5$ CD11b$^+$ monocytic BM cells that were isolated by biotinylated anti-CD11b (1:100; clone M1/70; ThermoFisher Scientific) and anti-biotin microbeads (Miltenyi Biotec) were seeded per well on 24-well plates in osteoclast differentiation medium as described above. The differentiation of MN-OCs and DC-OCs took 4–5 days and 5–6 days, respectively.

***Immunoblotting.*** Cells were washed with PBS and directly lysed by addition of boiling 2X Laemmli buffer containing phosphatase inhibitors (5 mM sodium orthovanadate, 20 mM sodium fluoride and 25mM β-glycerophosphate) 10 min at 95 °C [11]. Total lysates were separated on Bolt 8% polyacrylamide SDS gel (Novex) at 200 V constant then transferred to a nitrocellulose membrane (0,2 µm, GE Healthcare) in Bolt transfer buffer (Novex) for 1.5 hr at 115 mA constant. The membranes were blocked in Tris Buffered Saline (TBS, 50 mM Tris pH 7.2, 150 mM NaCl) with 3% BSA at room temperature for 1h with shaking, before being incubated overnight at 4°C with primary antibodies. Membranes were washed three times 5 min in TBS with 0.1% Tween 20 (TBS-T) and incubated for 1 hr at room temperature with HRP-coupled secondary anti-mouse (Sigma) or anti-rabbit antibodies (Cell Signaling) followed by three 5 min washes in TBS-T. The chemiluminescence signal was detected using Amersham ECL Prime Western Blotting Detection Reagent (GE Healthcare, Chicago, IL, USA) on the ChemiDoc Touch imaging system (Biorad). The intensity of each band is normalized to actin.

***RNA Extraction and qRT-PCR***. Total RNA was extracted at day 3, 5 or 7 of mOC differentiation using ready-to-use TRIzol Reagent (Ambion, Life Technologies, Austin, TX, USA) and purified with RNeasy Mini kit (Qiagen, Germantown, MD, USA). Complementary DNA was reverse transcribed from 1 µg total RNA with Moloney murine leukemia virus reverse transcriptase (Sigma-Aldrich) using dNTP (Promega, Madison, WI, USA) and random hexamer oligonucleotides (ThermoFisher, Waltham, MA, USA) for priming. qPCR was performed using SYBR green Supermix (OZYME, Saint-Cyr-l'École, France) in an ABI7500 Prism SDS real-time PCR detection system (Applied Biosystems, Foster City, CA, USA). The mRNA content was normalized to β-actin mRNA and quantified using the $2^{-\Delta\Delta Ct}$ method [12] Primers used for amplification of cDNA of NFACT-1, TRAP, Cathepsin K, DC-STAMP, β3-integrin and actin were purchased from Sigma-Aldrich and sequences are described in[12].

***Immunofluorescence and live imaging.*** Immunofluorescence experiments on glass coverslips or on bone slices were performed as described [13]. Briefly, cells were fixed with paraformaldehyde (PFA, 3.7%, Sigma-Aldrich), sucrose 30mM in PBS (Gibco), permeabilized with Triton X-100 0.3% (Sigma-Aldrich) for 10 min, and blocked with BSA (1% in PBS) for 30 min. Cells were incubated with primary antibodies for 1 hour, washed in PBS and then incubated with matching AlexaFluor secondary antibodies (2 µg/mL, Cell Signaling Technology), fluorescently labeled phalloidin (Invitrogen), and DAPI (500 ng/mL, Sigma Aldrich) for 30 min. After several washes with PBS, coverslips were mounted on a glass slide using fluorescence mounting medium (Dako) and stored at 4°C. The fusion index is defined as the number of nuclei present in a multinucleated giant cell (>2 nuclei) relative to the total number of nuclei per field [4,13]. Quantification of osteoclast fusion index, number of nuclei per multinucleated cells and area occupied by multinucleated cells was performed by using a semi-automatic quantification with a home-made ImageJ macro, allowing the study of more than 3,000 cells per condition. Thin and thick TNTs were identified by phalloidin and α-tubulin staining, and counted on at least 200 cells per condition. The number of sealing zones per surface, the area occupied by sealing zones, their circularity and thickness (3 zones per structures) were quantified after phalloidin staining of mature osteoclasts plated on bone slices. Most of the images were acquired using a Zeiss Axio Imager M2 and a 20X/0.8 Plan Apochromat or 40X/0.95 Plan Apochromat objectives (Zeiss). Images were acquired with a ORCA-flash 4.0 LT (Hamamatsu) camera and processed using the Zeiss Zen software. In some cases, super-resolution microscopy images (Figure 1A-C, Figure 4C and 4G, lower panels, Figure 5E, Figure S1B and Figure S4) were obtained with an Elyra 7 lattice SIM microscope and laser illumination sources (Zeiss). Images were processed with ImageJ and Photoshop software. Brightfield live imaging of hOCs was performed using a Zeiss LSM710 confocal microscope that uses a Zeiss AXIO Observer Z1 inverted microscope stand with transmitted (HAL), UV (HBO) and laser illumination sources or an IncuCyte ZOOM live cell imaging system (Essen Bioscience), in both cases acquiring images once per hour. For some live imaging experiments, we used 1:1 mixed cultures of control and KO mOCs transfected with Lifeact-mcherry or -GFP lentiviruses, respectively; as described [14].

***Bone resorption assays.*** To assess bone resorption activity, mature osteoclasts were detached using Accutase treatment (Gibco Technology, ThermoFischer Scientific, Courtaboeuf, France) 10 min, at 37°C, and cultured on bovine cortical bone slices (IDS Nordic Biosciences, Paris, France) for 24h in medium supplemented with M-CSF (25 ng/mL) and RANKL (100 ng/mL) at $1.10^5$ for hOCs. For mOCs, hoxB8 precursors were directly seeded on bovine cortical bone slices at a concentration of $5x10^4$cell/well (96-well/plate). Following complete cell removal by immersion in water and scraping, bone slices were stained with toluidine blue to detect resorption pits under a light microscope (Leica DMIRB, Leica Microsystems). The surface area of bone degradation was quantified manually with ImageJ software. Resorption pits and trenches were measured as described [15].

***Scanning Electron microscopy.*** hOCs and mOCs at day 3 of differentiation were fixed using 0.1 M sodium cacodylate buffer supplemented with 2.5% (v/v) glutaraldehyde. Cells were then washed three times 5min in 0.2 M cacodylate buffer (pH 7.4), post-fixed for 1h in 1% (wt/vol) osmium tetroxide in 0.2 M cacodylate buffer (pH 7.4), and washed with distilled water. Samples were dehydrated through a graded series (25 to 100%) of ethanol, transferred in acetone and subjected to critical point drying with $CO_2$ in a Leica EM CPD300. Dried specimens were sputter-coated with 3 nm platinum with a Leica EM MED020 evaporator and were examined and photographed with a FEI Quanta FEG250.

***Flow Cytometry.*** Cells were harvested with Accutase (Sigma-Aldrich), washed in PBS and collected by centrifugation at 500xg for 5 min, then stained with the LIVE/DEAD kit (Fisher Scientific) according to the manufacturer's instructions. Cells were counted and aliquoted into a 96-well plate at $3x10^5$ cells/well, labeled with 100 µL fluorescently conjugated β3-integrin antibodies for 30 min at 4°C, then washed twice in PBS. Data were acquired using a BD Fortessa X20 flow cytometer (BD Biosciences) driven by BD FACS Diva software, and analyzed using FlowJo (Tree Star, USA).

***Atomic force microscopy-based spectroscopy.*** CTL and moesin KO osteoclast precursors were plated into a 35mm Fluorodish (WPI) 24h before use. 30min prior to the AFM experiment, the serum concentration in the medium was reduced to 2% FBS. qp-SCONT cantilevers (Nanosensors) were mounted on a CellHesion 200 AFM (Bruker), connected into an Eclipse Ti inverted light microscope (Nikon). Cantilevers were calibrated using the contact-based approach, followed by coating with 4 mg/ml Concanavalin A (Sigma) for 1 hour at 37°C. Cantilevers were washed with 1xPBS before the measurements. MCA (Membrane-to-Cortex Attachment) was estimated using dynamic tether pulling: Approach velocity was set to 0.5 µm/s, with a contact force of 200 pN, and contact time was varied between 100 ms to 10 s, aiming at maximizing the probability to extrude single tethers. The cantilever was then retracted for 80 µm at a velocity of 2, 5, 10 or 30 µm/s. Tether force at the moment of tether breakage was recorded at a sampling rate of 2000 Hz. Resulting force curves were analyzed using the JPK Data Processing Software and the resulting force-velocity data was fitted to the Brochard-Wyart model [16] to allow estimation of an MCA parameter Alpha that is proportional to the density of binders (i.e. the active MCA molecules) and the emerging effective viscosity [17].

***Statistical analysis.*** All statistical analyses were performed using GraphPad Prism 9 (GraphPad Software Inc.). Two-tailed paired or unpaired t-tests were applied on data sets with a normal distribution (determined using Kolmogorov-Smirnov test), whereas two-tailed Mann-Whitney (unpaired test) or Wilcoxon matched-paired signed rank tests were used otherwise. When multiple comparisons were done, the statistical analyses used are detailed in the corresponding figure legend. $p<0.05$ was considered as the level of statistical significance (*, $p\leq0.05$; **, $p\leq0.01$; *** $p\leq0.001$; **** $p\leq0.0001$). n.s. not significant.

#### SUPPLEMENTAL FIGURE LEGENDS

**Supplemental Figure 1** (related to Figure 1). **A.** Experimental design of osteoclast differentiation from blood human CD14$^+$ monocytes (hOC, upper panel) or from a murine cell line immortalized thanks to the *HoxB8* gene (mOC, lower panel). **B.** Representative immunofluorescence analysis showing thin (white arrowheads) and thick tunneling nanotubes (TNTs) (orange arrowheads): F-actin (phalloidin, white), nuclei (DAPI, cyan) and microtubules ($\alpha$-tubulin, orange). Scale bar, 10 µm. **C.** Brightfield confocal images from a time-lapse movie of hOC fusion from protrusions emanating from the 2 fusing cells (upper panel) or at the tip of one TNT (lower panel) (hour:min). Dashed green and red lines delineate the nuclei before cell fusion and dashed orange lines after fusion. Arrowheads show TNT-like protrusions. Scale bar, 10 µm. See also **Movie 4&5**.

**Supplemental Figure 2** (related to Figure 2). **A-B.** Representative Western blot analysis of the level of ezrin, radixin, moesin during murine (A, mOC) and human (B, hOC) osteoclast differentiation, actin was used as loading control. **C**. Representative Western blot analysis of the level of ezrin, radixin, moesin and activated ERM (P-ERM) in the three individual KO mOC. **D.** Quantification of fusion index in control (CTL) and ezrin and radixin KO mOC, n=2 independent experiments, 3 images/experiment, 1000 nuclei/image, SD are shown. n.s. not significant. **E.** Quantification of the area occupied by osteoclasts in control (CTL) *versus* moesin KO (MKO) mOC after microscopy analysis, n=3 independent experiments, 4 images/experiment, 1000 nuclei/image. **F.** Flow cytometry analysis of the percentage of $\beta$3-integrin-positive cells in control (CTL) *versus* moesin KO (MKO) mOC. Each circle represents an independent experiment (n=6, SD are shown). **G.** Quantification of mRNA expression of genes overexpressed in osteoclasts measured by RT-PCR in control (CTL, blue) *versus* moesin KO mOC (orange) at day 3, 5 and 7 of differentiation. Actin mRNA level was used as control. SD are shown, n=3 independent experiments. **H -I**. Western blot analysis of moesin and activated ERM (P-ERM) expression level in hOC after treatment at day 0 with non-targeting siRNA (siCTL) or siRNA targeting moesin (siM), at day 3, 6 and 10 of differentiation. (H) Representative experiment and (I) quantification of moesin (left) and P-ERM (right) expression level normalized to actin. Each circle represents a single donor, SD are shown, n=7-8. Predicted molecular weight are indicated on Western Blots.

**Supplemental Figure 3** (related to Figure 2). **A-C.** Effect of lysophosphatidylcholine (LPC) treatment on hOC fusion and activated ERM expression level (P-ERM). At day 6 of differentiation, hOC were treated with LPC (+LPC, 17h), or treated and then washed (90 min) (+/- LPC) or no treated (control, CTL). (A-B) Microscopy analysis of hOC fusion. (A) Representative microscopy images: F-actin (phalloidin, white) and nuclei (DAPI, cyan). Scale bar, 50 µm. (B) Quantification of fusion index (each circle represents a single donor, n=3). (C) Western blot analysis of P-ERM expression level in each condition, normalized to actin. Representative blot (left panel) and quantification (right panel). Each circle represents a single donor, n=4. **D.** P-ERM signal by Western blot analysis in murine inflammatory osteoclasts (DC-OC) *versus* control osteoclasts (MN-OC). See material and methods. Representative blot of P-ERM expression level in each condition (upper panel) and quantification of P-ERM expression level, normalized to actin (lower panel). Each circle represents a single experiment (n=4, SD are shown). **E-H.** Effect of HIV-1 infection on fusion index and on P-ERM signal in human osteoclasts (E-F) and macrophages (G-H). (E-F) At day 6 of differentiation, hOC were infected with the viral strain NLAD8-VSVG (+HIV-1) or not (CTL) and analyzed 8 days post-infection. (E) Quantification of the fusion index and (F) representative Western blot analysis of P-ERM expression level (left), and quantification of P-ERM expression level, normalized to actin (right, each circle represents a single donor, n=4). (G-H) At day 6 of differentiation, macrophages were infected with the viral strain NLAD8-VSVG (+HIV-1) or not (CTL) and analyzed 7 days post-infection. (G) Quantification of the fusion index and (H) representative Western blot analysis of P-ERM expression level (left) and quantification of P-ERM expression level, normalized to actin (right). Each circle represents a single donor (n=5-6, SD are shown). Predicted molecular weight are indicated on Western Blots.

**Supplemental Figure 4** (related to Figure3). Super-resolution microscopy images showing moesin localization in hOC on glass coverslides (A-C, Scale bar, 10 µm) or on bone (D, Scale bar, 5 µm): F-actin (phalloidin, magenta), nuclei (DAPI, cyan) and moesin (green). F-actin structures: (A) TNTs, (B) zipper-like structures, (C) a podosome belt and (D) a sealing zone.

**Supplemental Figure 5** (related to Figure3). **A.** Representative immunofluorescence images showing activated ERM (P-ERM) at TNTs. F-actin (phalloidin, magenta), nuclei (DAPI, cyan) and P-ERM (green). Scale bar, 10 µm. See also **Movie 7**. **B.** Representative microscopy images (F-actin, phalloidin, grey) illustrating the increase of TNT number after moesin depletion: (left) Moesin KO *versus* CTL mOC and (right) siMoesin *versus* siCTL hOC. Scale bars, (left) 100 µm and (right) 50 µm. **C.** Representative image of a 1:1 mixed culture of mOC control (CTL, transduced with mcherry-lifeact) and moesin KO (transduced with GFP-lifeact) seeded on glass coverslides at day 3 of differentiation. Arrowheads show TNT-like protrusions. Scale bar, 20 µm.

**Supplemental Figure 6** (related to Figure 4). **A.** Representative image from a culture with 1:1 ratio of mOC control (CTL, transduced with mcherry-lifeact) and moesin KO (transduced with GFP-lifeact) seeded on glass coverslips at day 4 of differentiation. Arrowheads show green podosome belts. Scale bar, 20 µm. **B.** Effect of moesin KO on sealing zone formation in mOC.

mOC control (CTL) *versus* moesin KO (MKO) were cultured for 5 days on glass coverslips, detached and then seeded for additional 2 days on bone slices. Quantification of the area occupied by sealing zones (left) and circularity of sealing zones (right) (=3 independent experiments, 2 bone slices/condition, SD are shown). n.s. not significant. **C.** Effect of moesin depletion on sealing zone formation in hOC. 6 day-differentiated hOC on glass coverslips treated at day 0 with siCTL or siMoesin were detached and seeded for additional 24h on bone slices. Quantification of the number of nuclei per cells forming sealing zones (n=4 donors, 2 bone slices/condition).

**Supplemental Figure 7** (related to Figure 4). **A.** Experimental design of moesin depletion in late stages of hOC differentiation. **B.** Effect of moesin depletion (siMoesin) on hOC fusion: quantification of fusion index after microscopy analysis of 10 day-differentiated hOCs on glass coverslips treated at day 6 with siCTL or siMoesin. Each circle represents a single donor (n=7, SD are shown). **C.** Western blot analysis of moesin and P-ERM expression levels after moesin depletion in late stages of hOC differentiation. Representative Western blot analysis (left) and quantification of moesin and P-ERM signals (right), normalized to actin. Predicted molecular weight are indicated. Each circle represents a single donor (n=8, SD are shown). **D-H.** Effect of moesin depletion (siMoesin) in mature hOC on bone degradation (D), morphology of the resorbed area (E) and sealing zone (SZ) formation (F-H). 10 day-differentiated hOCs on glass coverslips treated at day 6 with siCTL or siMoesin were detached and seeded for additional 24h on bone slices. (D) Representative images of bone degradation (left, scale bar, 50 µm) and quantification of bone eroded surface (%) using semi-automatic quantification (right) (n=4 donors, 2 bones slices/condition). (E) Quantification of the percentage of trenches (n=2 independent experiments, 2 bone slices/condition, SD are shown). (F) Representative microscopy images of sealing zone visualized by F-actin staining (phalloidin, white, scale bars, 20 µm); and (G) quantification of the number of sealing zones (number of SZ per bone surface (right) and the percentage of area covered by SZ (left) (n=6 donors, 2 bone slices/condition). (H) Effect of moesin depletion (siMoesin) in mature hOCs on SZ thickness and organization. (Left) Representative microscopy images of sealing zones visualized by F-actin and vinculin stainings (phalloidin in pink and vinculin in green). Scale bars, 10 µm. (Right) Signal intensity profile of vinculin (green) and actin (green) along the white lines (left) in siCTL (upper panel) or siMoesin (lower panel) (n=1 experiment, n=5-6 cells/conditions). n.s. not significant.

**Supplemental Figure 8** (related to Figure 5). **A.** Effect of calyculin and staurosporine treatment on ERM activation (P-ERM), used as positive and negative control for Western blot analysis of ERM activation, respectively. 6 day-hOCs were treated or not (CTL) with calyculin and staurosporine. Representative Western blot analysis (left) and quantification of P-ERM signal normalized to actin (right). Each circle represents a single donor (n=6, SD are shown). **B.** Effect of NSC23 and ML 141 on ERM activation. 6 day-hOCs were treated or not (CTL) with NSC23 and ML 141, drugs targeting Rac1/2 and Cdc42 respectively. Representative Western blot analysis (left) and quantification of P-ERM signal normalized to actin (right). Each circle represents a single donor (n=5, SD are shown). **C.** Effect of Y27632 (ROCK kinase inhibitor) treatment on P-ERM signal. Representative Western blot analysis (left) and quantification of P-ERM signal normalized to actin (right). Each circle represents a single donor (n=4, SD are shown). Predicted molecular weight are indicated on Western Blots. n.s. not significant.

**Supplemental Figure 9** (related to Figure 6)

**A.** Representative immunofluorescence images of histological analysis of femurs from WT mice: nuclei (DAPI, blue) and moesin (section 1, green) or cathepsin K (section 2, green). Scale bar, 20 µm. Sections 1&2 are serial sections. White arrowheads show osteoclasts**. B.** Representative X-Ray images of the whole skeleton of WT and *Msn*$^{-/-}$ mice. **C.** Histograms indicate means ± SD of cortical bone parameters analyzed by microcomputed tomography. Animal groups were composed of 6 mice of each genotype, and 11 femora were analyzed in total for *Msn*$^{-/-}$ mice and 9 femora for the WT mice group. n.s. not significant.

**SUPPLEMENTAL MOVIES**

**Supplemental Movie 1** (related to Figure 1C, left panel) – Z-stack reconstitution of super-resolution microscopy images showing TNTs in hOCs with a colored-coded Z of F-actin signal (phalloidin).

**Supplemental Movie 2** (related to Figure 1C, right panel) - Z-stack reconstitution of super-resolution microscopy images showing TNTs in mOCs with a colored-coded Z of F-actin signal (phalloidin).

**Supplemental Movie 3** (related to Figure 1D) - Time-lapse of microscopy images (DIC from confocal microscopy) showing the fusion of hOCs. 1 image every 5 min.

**Supplemental Movie 4** (related to Figure S1C, upper panel) - Time-lapse of microscopy images (DIC from confocal microscopy) showing the fusion of hOCs. 1 image every 5 min.

**Supplemental Movie 5** (related to Figure S1C, lower panel) - Time-lapse of microscopy images (DIC from confocal microscopy) showing the fusion of hOCs. 1 image every 5 min.

**Supplemental Movie 6** (related to Figure 2C) - Time-lapse of microscopy images (DIC from Incucyte®) showing the differentiation and fusion into giant cells of control (CTL, left) and moesin KO (right) mOCs. 1 image every 1 hour.

**Supplemental Movie 7** (related to Figure S5Alower panel) – 3D reconstitution of confocal microscopy images showing activated ERM (P-ERM) at TNTs. F-actin (phalloidin, magenta), nuclei (DAPI, cyan) and P-ERM (green).

#### Supplemental Figure 1 - related to Figure 1

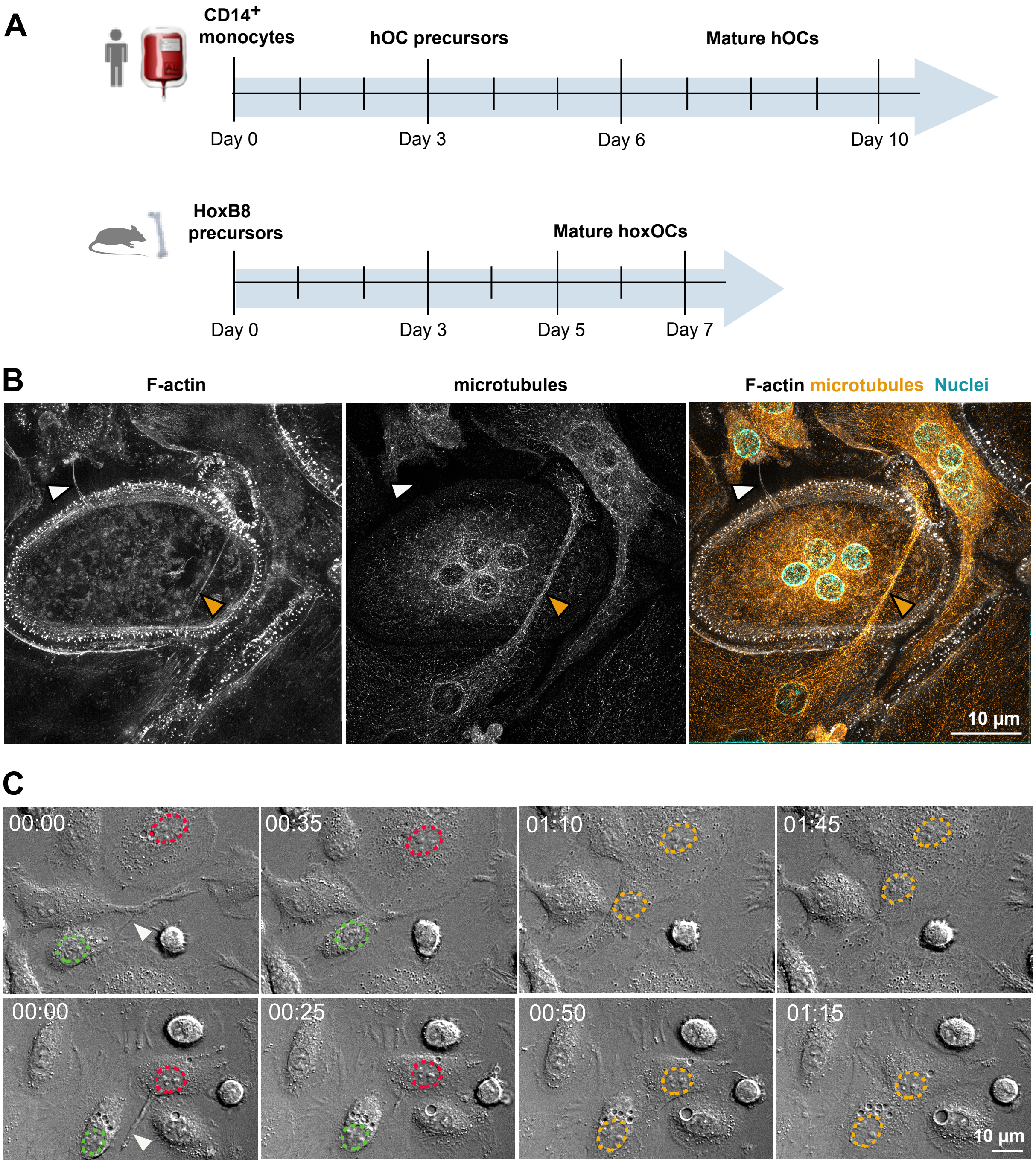

#### Supplemental Figure 2 - related to Figure 2

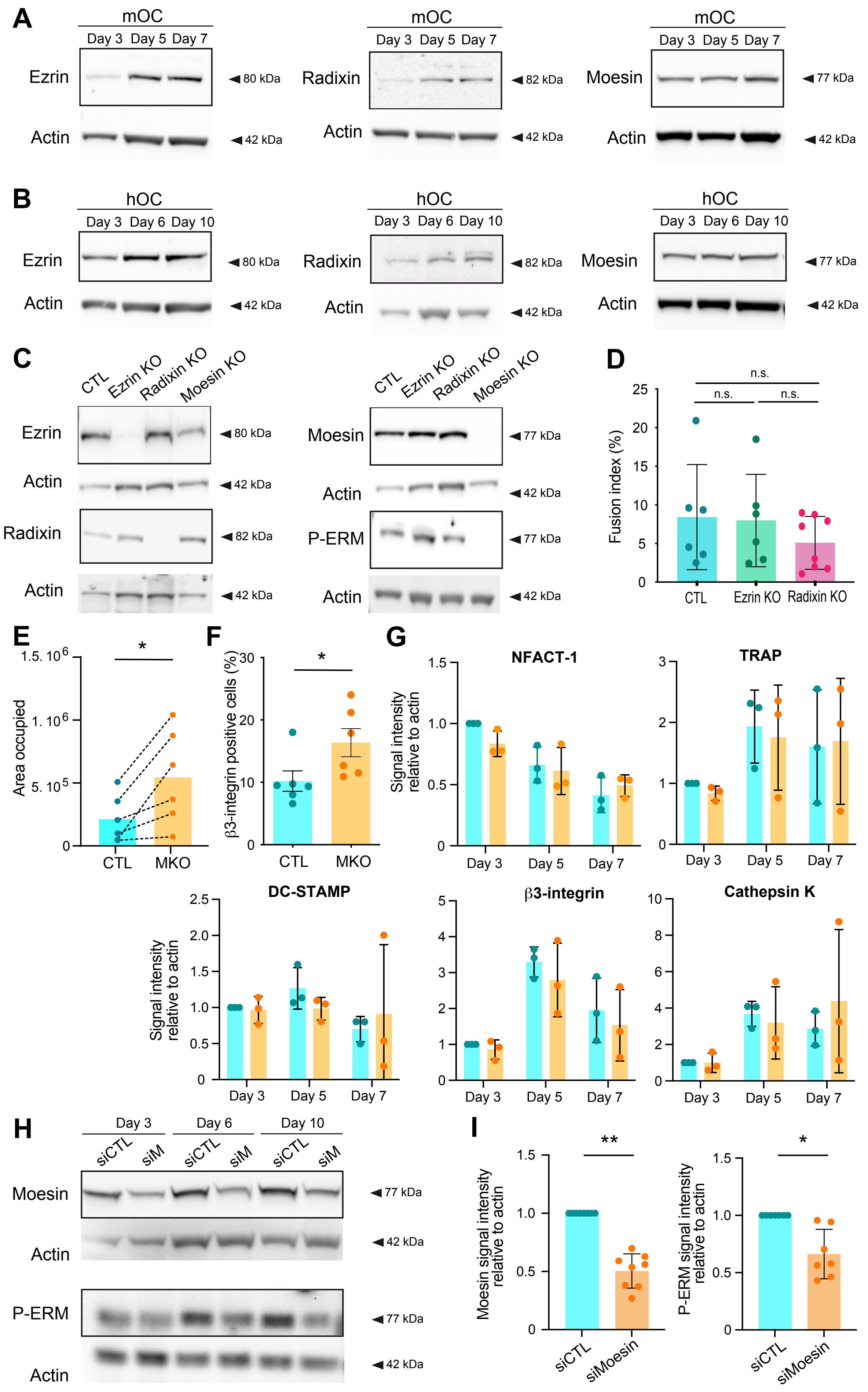

#### Supplemental Figure 3- related to Figure 2

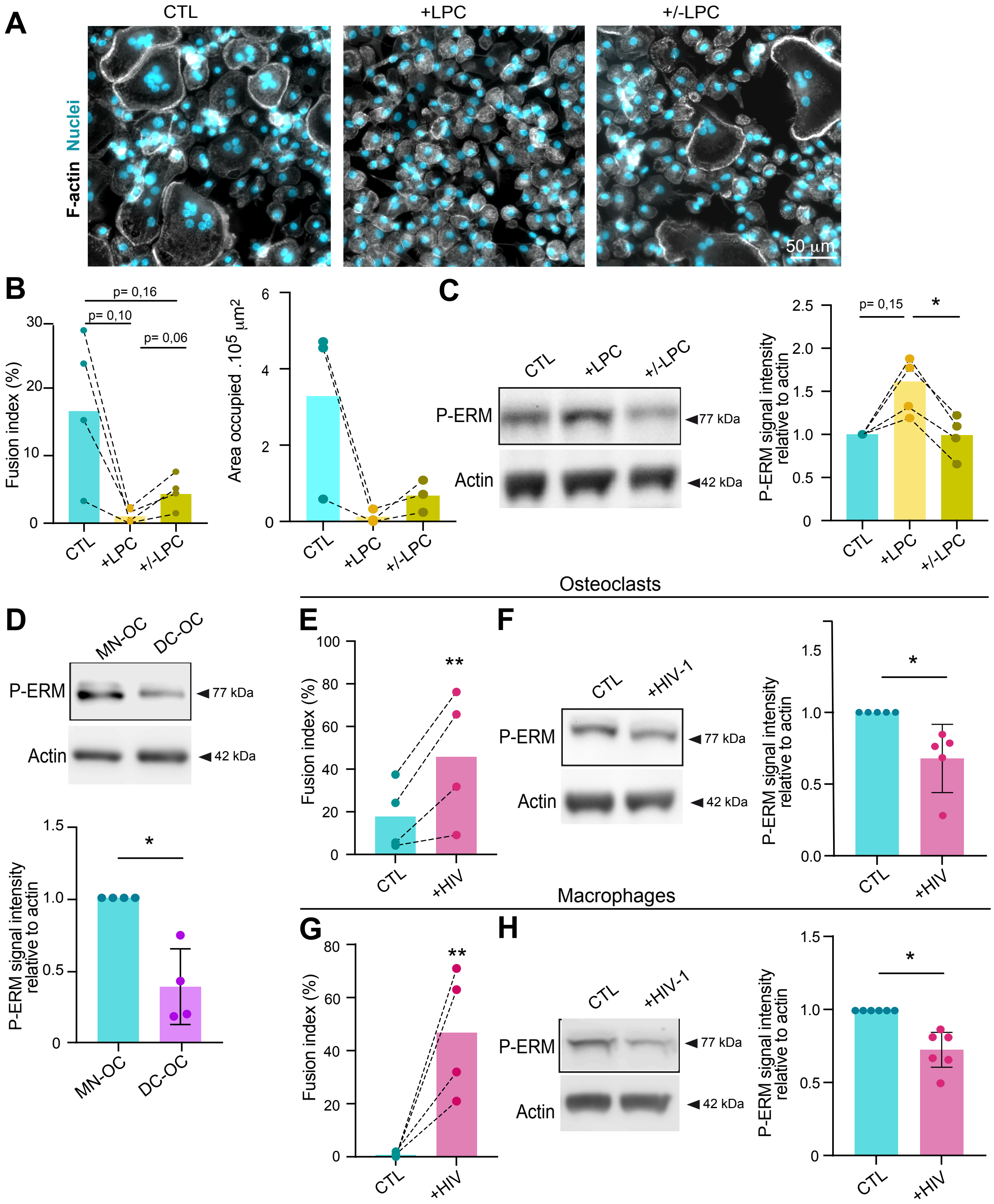

#### Supplemental Figure 4 - related to Figure 3

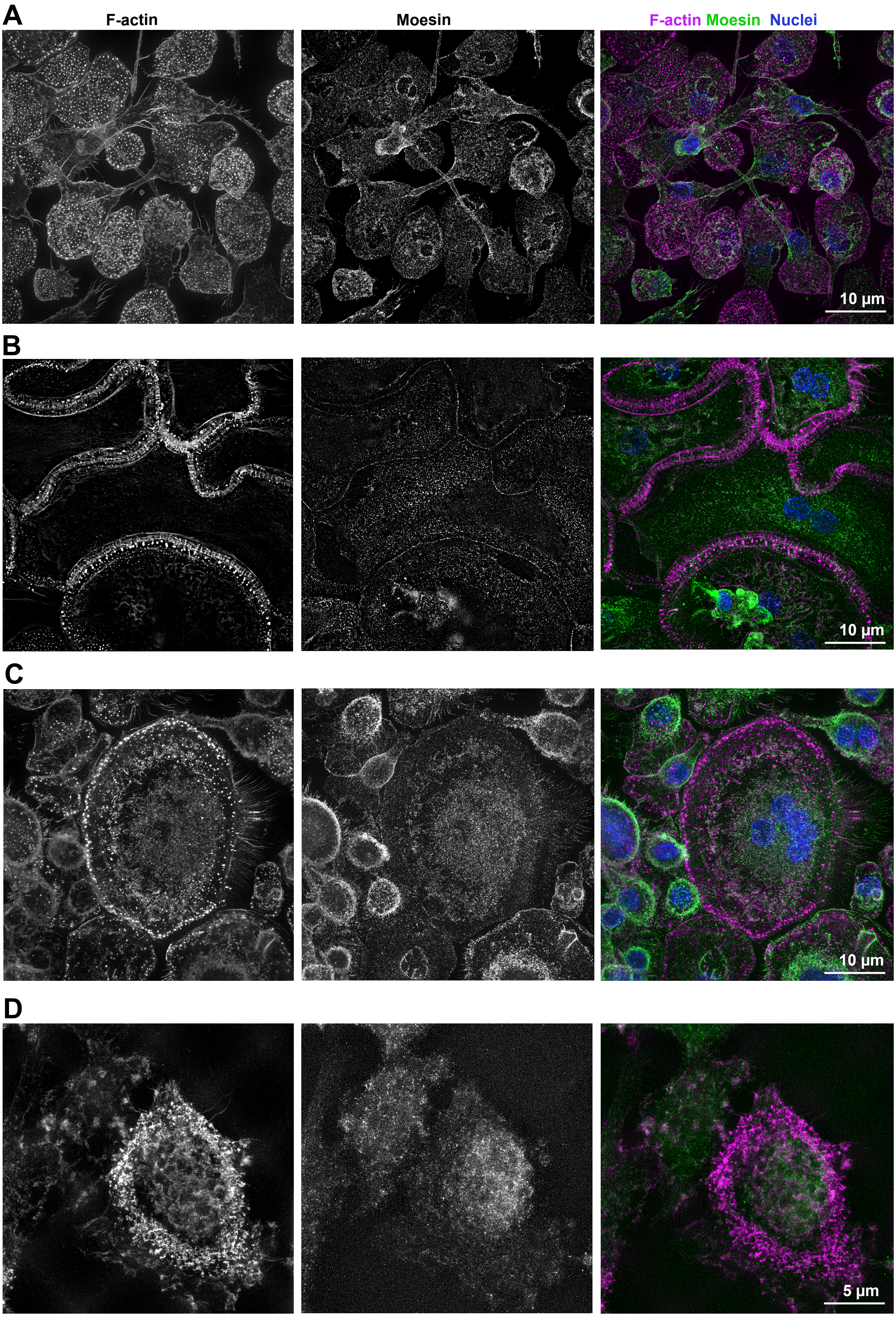

**Supplemental Figure 5 - related to Figure 3**

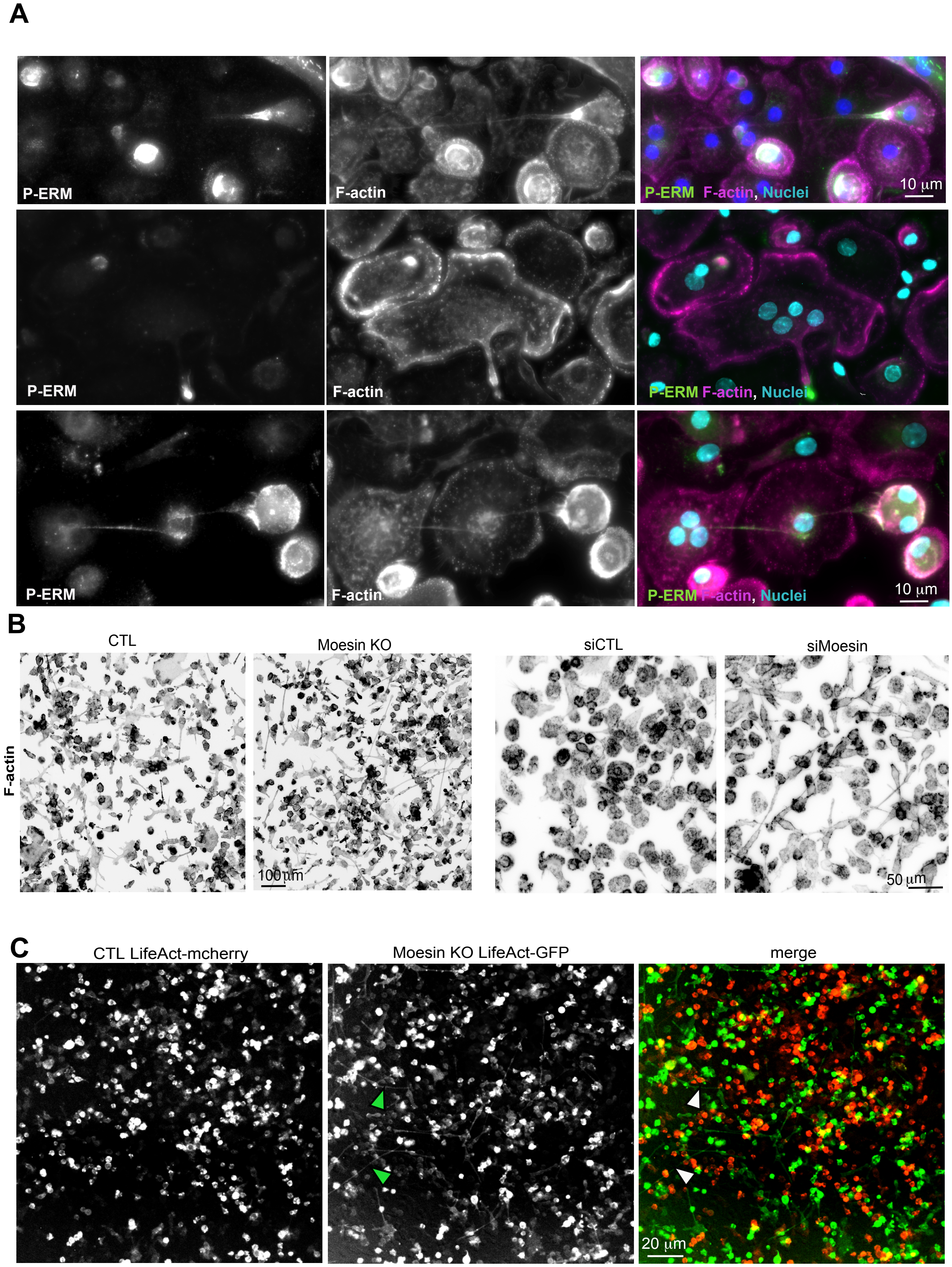

**Supplemental Figure 6 - related to Figure 4**

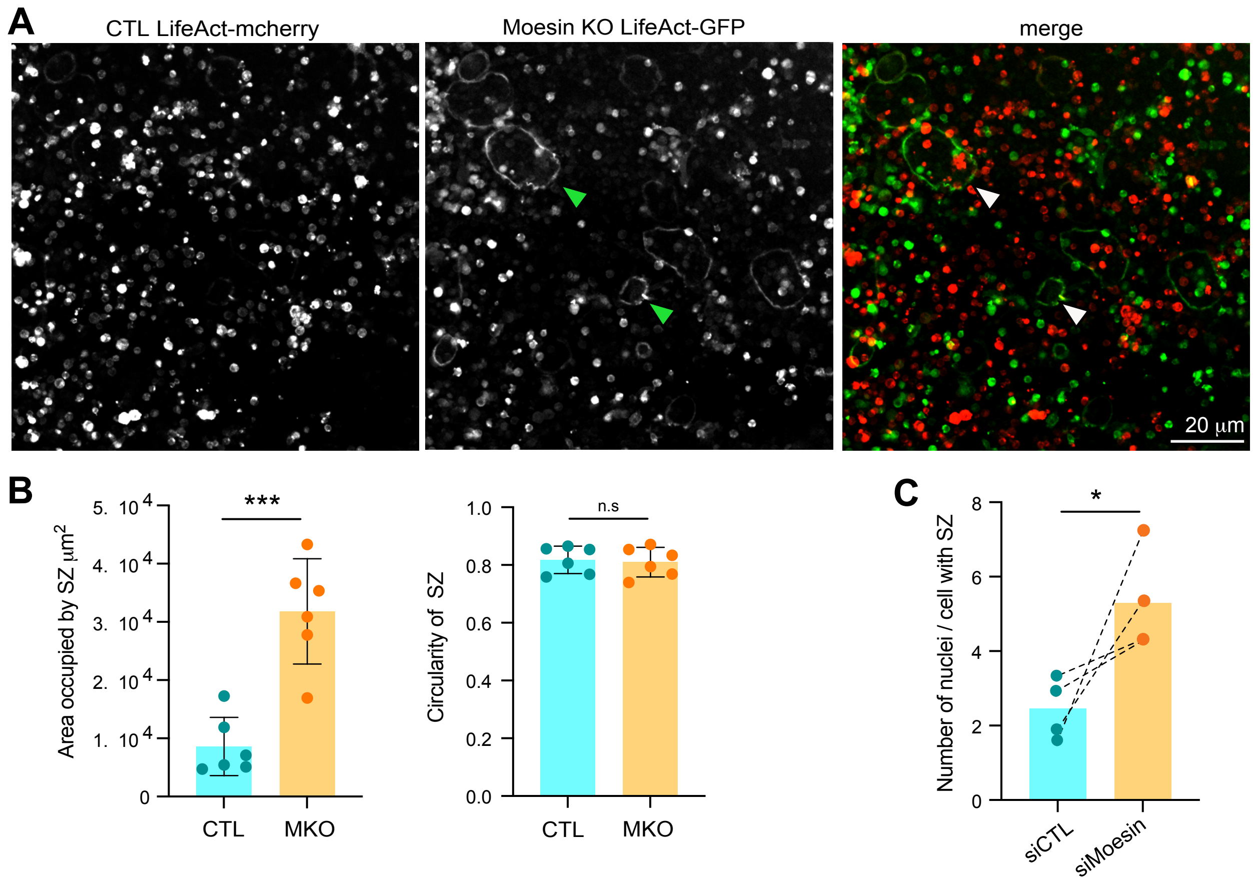

### Supplemental Figure 7 - related to Figure 4

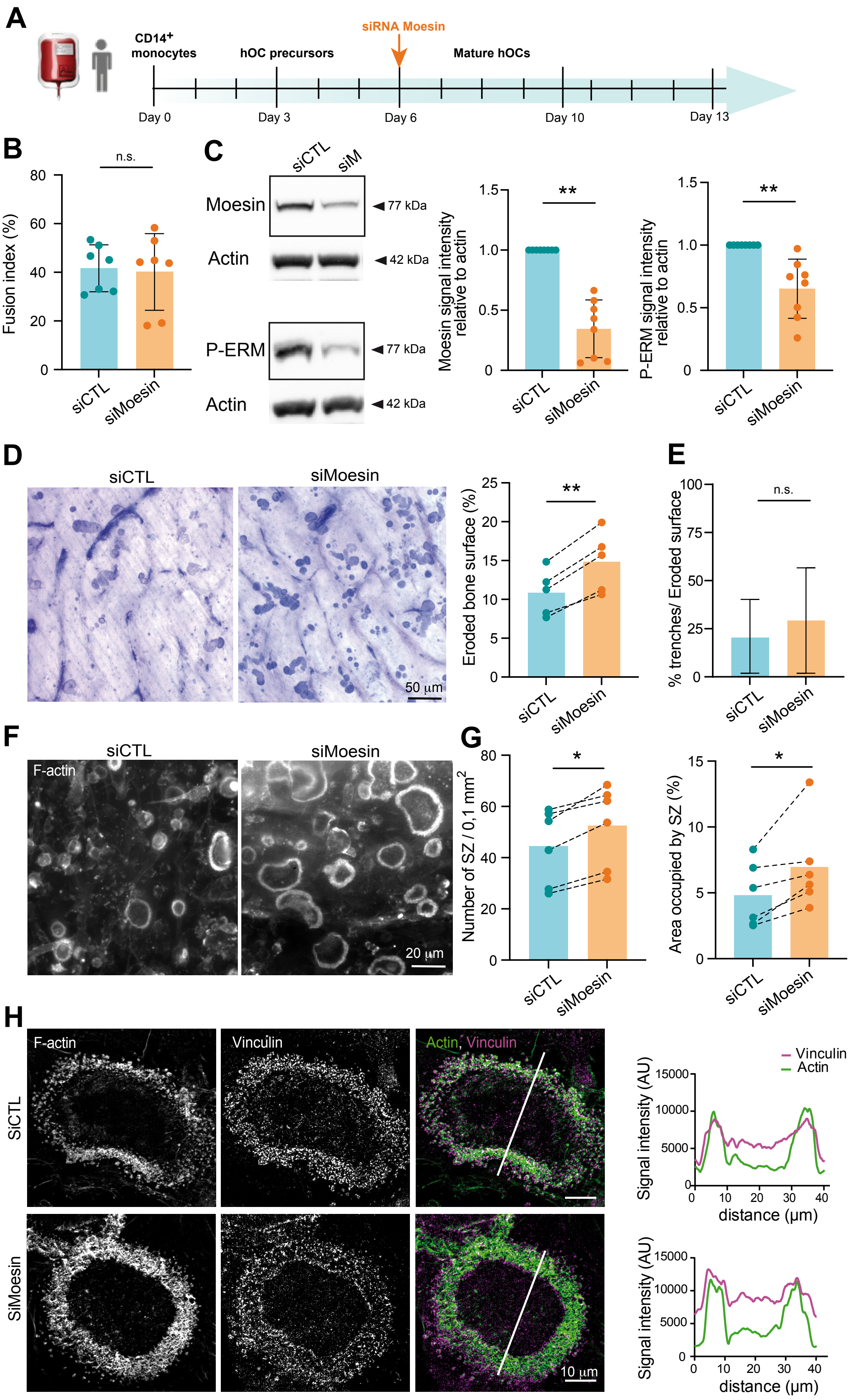

#### Supplemental Figure 8 - related to Figure 5

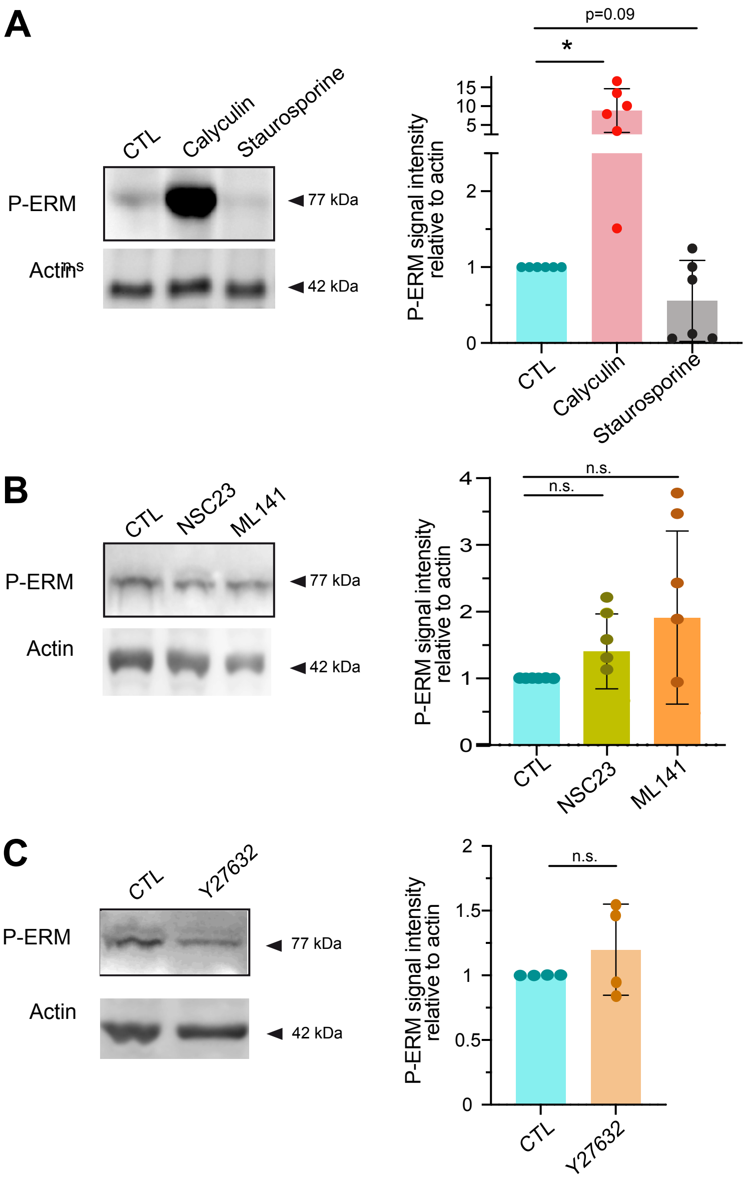

#### Supplemental Figure 9 - related to Figure 6

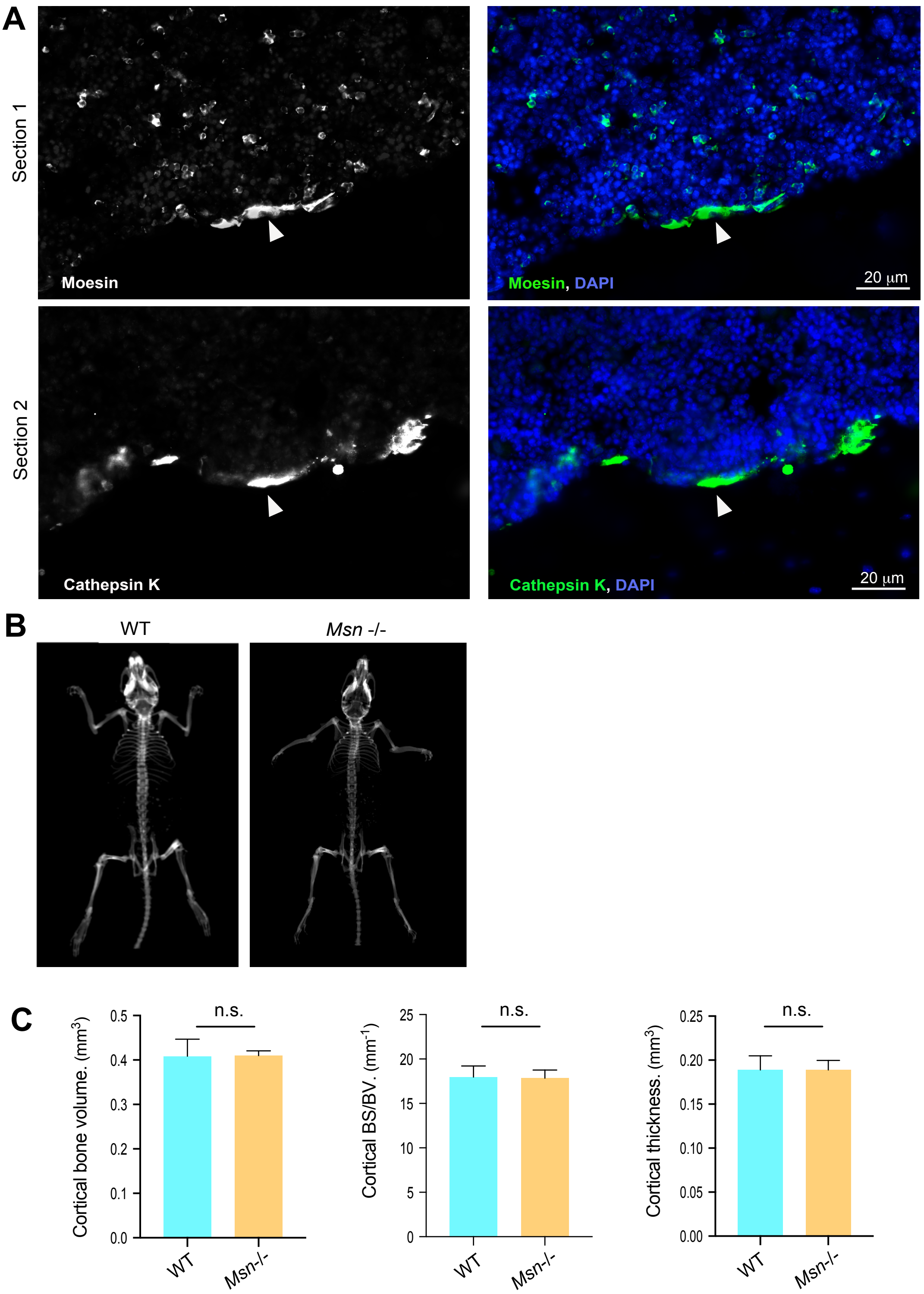
